# Regionalization of the nervous system requires axial allocation prior to neural lineage commitment

**DOI:** 10.1101/229203

**Authors:** Vicki Metzis, Sebastian Steinhauser, Edvinas Pakanavicius, Mina Gouti, Despina Stamataki, Robin Lovell-Badge, Nicholas M Luscombe, James Briscoe

## Abstract

**Summary:** Neural induction in vertebrates generates a central nervous system that extends the rostral-caudal length of the body. The prevailing view is that neural cells are initially induced with anterior (forebrain) identity, with caudalising signals then converting a proportion to posterior fates (spinal cord). To test this model, we used chromatin accessibility assays to define how cells adopt region-specific neural fates. Together with genetic and biochemical perturbations this identified a developmental time window in which genome-wide chromatin remodeling events preconfigure epiblast cells for neural induction. Contrary to the established model, this revealed that cells commit to a regional identity before acquiring neural identity. This “primary regionalization” allocates cells to anterior or posterior regions of the nervous system, explaining how cranial and spinal neurons are generated at appropriate axial positions. These findings prompt a revision to models of neural induction and support the proposed dual evolutionary origin of the vertebrate central nervous system.

## Introduction

The acquisition of neural identity, known as neural induction (Stern, 2006) represents one of the most widely studied events in embryogenesis. In vertebrates, this process begins at gastrulation and continues as the principal axis elongates resulting in a nervous system extending along the anterior-posterior (AP) length of the body. The critical role of the organizer in specifying neural fate from ectoderm was initially established by the pioneering work of Spemann and Mangold (Spemann and Mangold, 1924). Attention then turned to identifying the inducing signals emanating from the organizer and understanding how different rostral-caudal regions of the nervous system are generated (Anderson and Stern, 2016; Stern, 2001; Stern et al., 2006).

Several models have been proposed to explain rostral-caudal regionalisation. Embryological experiments led Otto Mangold to suggest separate activities are responsible for inducing distinct regions of the nervous system (Mangold, 1933). He proposed that different parts of the organiser, or the organiser at different times, produced these distinct signals. Subsequently, Nieuwkoop, building on the work of Conrad Waddington (Waddington, 1940), proposed a two-step mechanism to explain the formation and regionalization of the nervous system known as “activation-transformation” (Nieuwkoop, 1952). This hypothesis contends that cells first adopt a neural identity equivalent to the anterior nervous system (“activation”). “Transformation”, in a subsequent step, is responsible for converting a proportion of these cranial-like cells to more caudal fates such as the midbrain, hindbrain and eventually, spinal cord (Nieuwkoop and Nigtevecht, 1954; Stern, 2001).

In this view, anterior neural cells are considered the precursors of the entire nervous system. This implies that cells that form the nervous system are first specified with an anterior identity before they acquire more caudal axial fates such as hindbrain or spinal cord. Whether this mechanism is valid and applicable to all axial levels of the nervous system is unresolved. Nevertheless, it remains the prevailing view of nervous system regionalisation (Stern, 2001, 2005, 2006) and has influenced the development of methods for the directed differentiation of embryonic stem cells to specific classes of neurons, where regionalising signals are assumed to act after neural identity has been established in cells (Davis-Dusenbery et al., 2014; Wichterle et al., 2002).

The anterior nervous system in vertebrates, comprising fore-, mid- and hindbrain, has an anatomically and molecularly distinct origin from the spinal cord. The anterior nervous system is formed during gastrulation from cells that remain in the anterior epiblast. By contrast, spinal cord cells are produced during axis elongation by axial stem cells, often referred to as neuromesodermal progenitors (NMPs) (Henrique et al., 2015). These bipotent cells arise in the caudal lateral epiblast, adjacent to the node, and contribute progeny to both the paraxial mesodermal tissue and spinal cord (Garriock et al., 2015; Tzouanacou et al., 2009; Wymeersch et al., 2016). NMPs are exposed to WNT and FGF signalling and are marked by the expression of transcription factors *Sox2, T/Brachyury* and *Cdx1, 2, 4* (Gouti et al., 2014, 2017; Henrique et al., 2015; Tsakiridis et al., 2014; Wymeersch et al., 2016). Deletion of *T/Bra, Cdx* genes or the absence of WNT signalling severely abrogates axis elongation in mouse embryos, resulting in a failure to form spinal cord and somites at post occipital levels (Amin et al., 2016; Chawengsaksophak et al., 1997; Takada et al., 1994; Yamaguchi et al., 1999; Young et al., 2009). Thus, anterior and posterior parts of the nervous system are populated by distinct groups of cells. Similar to the case *in vivo*, timely pulses of WNT and FGF signals to ESCs that have acquired an epiblast-like state results in the generation of cells resembling NMPs found in embryos (Gouti et al., 2017; Koch et al., 2017). These cells have the capacity to differentiate into spinal cord progenitors that express 5’ *Hox* genes characteristic of thoracic and lumbar spinal cord (Gouti et al., 2014; Lippmann et al., 2015). Single-cell transcriptome analysis further emphasises the correspondence between in vitro and in vivo cell populations (Gouti et al., 2017; Koch et al., 2017). By contrast, ESCs that are differentiated to an epiblast state in the absence of a WNT pulse will generate neural progenitors that display a caudal limit at the level of the hindbrain and cervical spinal cord (Gouti et al., 2014; Lippmann et al., 2015). These observations appear to challenge the Activation-Transformation hypothesis, and are reminiscent of Mangold’s model of distinct mechanisms specifying different regions of the nervous system (Mangold, 1933).

To address the sequence of events that lead to the establishment of a regionalised nervous system an unbiased definition of neural cell identity is required. Early embryological experiments relied on morphological criteria to define cell types and thus do not provide sufficient molecular detail to understand nervous system regionalisation. More recently, gene expression has been used as a proxy for identity. However, this has raised further questions. While common gene regulatory networks (GRNs) are used to globally define neural progenitor (NP) populations, it remains a challenge to understand how these networks are established genome wide, leaving open the question of how neural progenitors become refined into functionally distinct neural cell types along the AP axis. For instance, the SoxB1 family of transcription factors play critical roles in neural progenitors along the AP axis and are broadly expressed, yet it remains unclear how they act at different axial levels (Kondoh et al., 2016). By contrast, distinct enhancer usage in cells has been used to define different cell types (Soucie et al., 2016) and has been shown to better resolve cell identity than conventional transcriptome comparisons (Corces et al., 2016). A repertoire of enhancers is known to drive AP-specific expression of genes that are broadly expressed throughout the nervous system, including the major neural regulators SOX2 and SHH (Epstein et al., 1999; Kutejova et al., 2016; Peterson et al., 2012; Uchikawa et al., 2003). Thus, regulatory element usage provides a reliable and objective correlate of cell identity (Buecker and Wysocka, 2012). One way to assay this is to systematically map and compare chromatin accessibility in different cell types. Techniques such as DNase-seq (Song and Crawford, 2010) and ATAC-seq (Buenrostro et al., 2013) provide this opportunity and the regions identified by these approaches show a high degree of correlation with active histone marks and known enhancers (Buenrostro et al., 2013; Lavin et al., 2014; Wu et al., 2016).

Here, we define the enhancer landscape using ATAC-seq, in cells with anterior (forebrain/midbrain), hindbrain and spinal cord identity. We take advantage of the temporal resolution afforded by the *in vitro* differentiation of ESCs into defined neural fates to determine how chromatin accessibility changes in time and how this relates to the progression of cells to anterior and posterior neural fates. Combined with *in vivo* validation, we show that the difference between anterior and posterior neural progenitors is reflected in their respective chromatin accessibility profiles. We provide evidence that AP identity precedes the acquisition of neural fate. Furthermore, we find that the genomic landscape of NMPs is distinct from other cell types and is dependent on the presence of CDX TFs that remodel the chromatin landscape in response to FGF and WNT signalling. This transition is essential not only to elicit induction of posterior *Hox* genes, but also to repress cranial neural fates. The ability to induce an NMP state in cells is transient (Gouti et al., 2014; Turner et al., 2014) and restricted to stages prior to the acquisition of neural identity; continual changes in the genomic accessibility of cells undergoing neural induction are sufficient to change the intrinsic response of a cell to the same extrinsic signals and the resulting cell fate identity. Together with genetic perturbations and alterations in the timing of posteriorizing signals, the data reveal that, contrary to the activation-transformation hypothesis, axial identity is established before neural induction. These findings are consistent with the proposed dual origin of the central nervous system during animal evolution (Arendt et al., 2016) and prompts a revision to models of neural induction and nervous system regionalization.

## Results

### In vitro generation of anterior, hindbrain or spinal cord neural progenitors

To define the sequence of events that commit neural cells to different anterior-posterior (AP) identities, we took advantage of mouse embryonic stem cells (ESCs), which, as shown previously, can be directed to differentiate into NPs with anterior, hindbrain or spinal cord identities (Gouti et al., 2014, 2017) (see Methods). In each case, ESCs were transferred from pluripotent conditions to serum-free media containing bFGF (FGF) to induce an epiblast identity. For anterior NPs, FGF was removed after three days and the SHH agonist SAG added, promoting ventral neural identity. By Day (D) 5 these cells expressed a mixture of forebrain and midbrain markers (Gouti et al., 2014). For the generation of hindbrain NPs, cells were exposed to retinoic acid (RA), in addition to SHH signals from D3 to D5 (Figure 1A). This produced visceral motor neuron (MN) progenitors expressing PHOX2B and somatic MNs expressing OLIG2, similar to the brainstem (Figure 1B) (Gouti et al., 2014; Pattyn et al., 1999, 2000). For the generation of spinal cord NPs, ESCs were cultured in the same serum-free media containing bFGF for two days (Figure 1A) and then received a 24 hour pulse of both FGF and WNT (FGF/WNT) signals from D2-3 (WNT signalling was induced with the GSK3β inhibitor, CHIR99021; see Methods). At D3, cells were transferred to medium containing RA and SHH signals, similar to the hindbrain condition (Figure 1A). At D5, this resulted in the generation of Olig2 positive spinal somatic MN progenitors (Figure 1B) but no visceral MNs (Figure 1B) and expressed *Hox* genes characteristic of cervical/brachial and thoracic regions (see Figure 3E-M).

**Figure 1.**
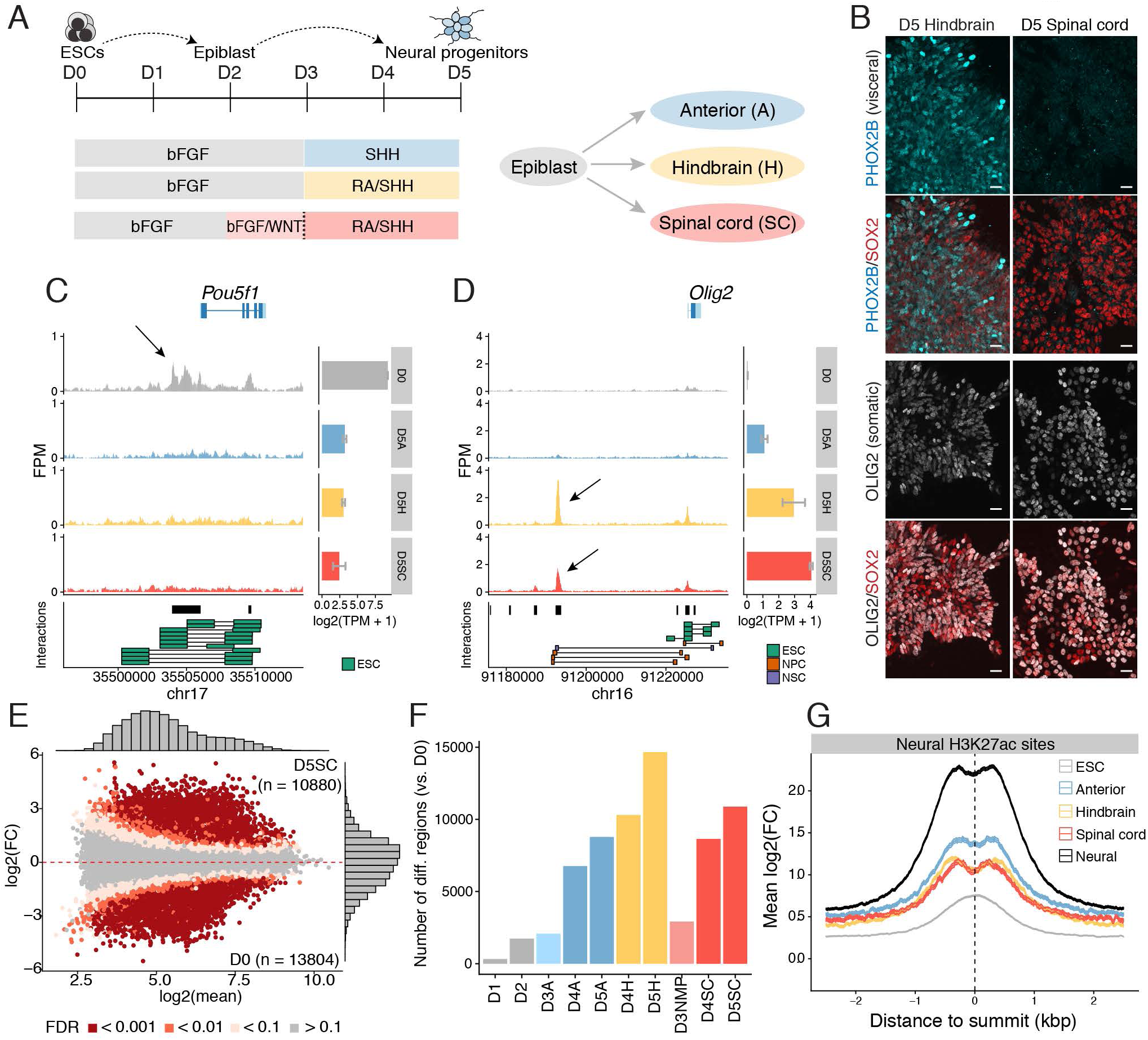
Regulatory element usage distinguishes cell states during neural induction. **(A)** Schematic of the five-day directed differentiation of mouse embryonic stem cells (ESCs) to neural progenitors that yields anterior (top), hindbrain (middle) or spinal cord (bottom) identities. Note that spinal cord progenitors are generated via an NMP state induced by the addition of FGF and WNT signals from day 2-3 (light pink shading). **(B)** Immunofluorescence on day 5 reveals that hindbrain progenitors generate a mixture of PHOX2B expressing visceral- and OLIG2 expressing somatic motor neuron progenitors. Spinal cord progenitors lack visceral- but generate OLIG2 expressing somatic motor neuron progenitors. Scale bars = 20 microns. **(C-D)** Genome browser tracks of ATAC-seq accessible regions (mm10 assembly) present in ESCs (day 0, grey) compared to day 5 anterior (blue), hindbrain (yellow) and spinal cord (red) progenitors, and associated gene expression levels determined by mRNA-seq (Gouti et al., 2014) for each stage as indicated on the right (Error bars = SEM). Cis-interactions indicated below represent known genomic interactions from published data (see Methods). ESCs show accessibility at *Pou5f1/Oct4* enhancers (C, arrow) unlike neural progenitors which repress *Oct4*. Instead open regions flank neural genes such as *Olig2* (D, arrow). (**E**) Genome wide comparison in accessibility between Day 5 spinal cord (D5SC) and Day 0 ESCs reveals differences in regulatory element usage (FDR?0.01). (**F**) The proportion of differential sites present in each condition compared to ESCs demonstrates the gradual change in accessibility during neural progenitor differentiation. (**G**) Both neural and AP-specific sites, but not ESC sites, are enriched in H3K27ac marks from neural progenitors (Peterson et al., 2012). bFGF= basic fibroblast growth factor; ESC=embryonic stem cell; D=day; FC=fold change; kbp=kilobase pairs; RA=retinoic acid; SHH=sonic hedgehog; TPM=transcripts per million.

**Figure 3.**
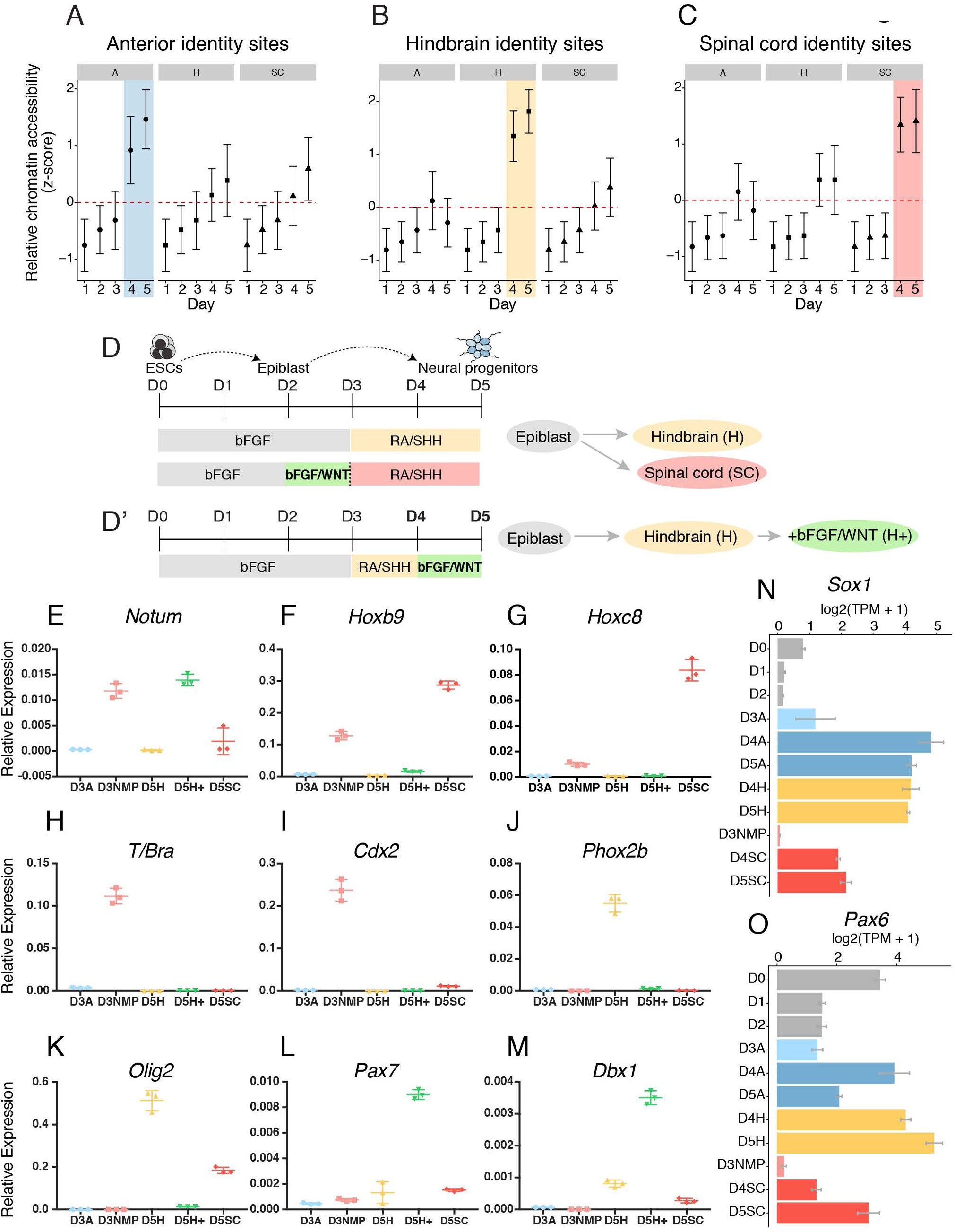
Axial identity is established in cells prior to neural identity. **(A-C)** The average accessibility (z-score) of region-specific sites over time in anterior (A), hindbrain (H) or spinal cord (SC) conditions. In each case (A-C), AP-specific sites become accessible between D3-4, the time point corresponding to the addition of neuralising signals to the culture medium. Spinal cord progenitors do not transiently open sites corresponding to anterior (A) or hindbrain (B) identity before opening spinal cord-specific sites (C). Error bars=SD. **(D)** Schematic of the differentiation used to generate hindbrain and spinal cord cells. The only difference between these two conditions is the addition of WNT signals between day 2-3 in the spinal cord condition, which is provided together with bFGF (bFGF/WNT). By contrast, hindbrain cells are only exposed to bFGF at this time (shaded in grey). **(D’)** Schematic of the differentiation used to test the posteriorising effect of bFGF/WNT in hindbrain neural progenitors. bFGF/WNT signals are provided to cells from day 4-5 (H+ condition, D’). **(E-M)** RT-qPCR data showing the relative expression of genes at D3 and D5 following the differentiation of cells to hindbrain, spinal cord or “hindbrain+” identity. The induction of the WNT signaling target gene *Notum* (E) is observed both at day 3 (D3NMP, following day 2-3 treatment with bFGF/WNT) and at day 5 (D5H+, following treatment with bFGF/WNT between day 4-5). By contrast, induction of posterior spinal cord *Hox* genes *Hoxb9* and *Hoxc8* is dependent on timing: induction at day 3 in D3NMP cells is observed following day 2-3 treatment with bFGF/WNT signals, but not at day 5 in D5H+ cells following day 4-5 treatment with the same signals (F-G). Similarly, the induction of *T/Bra* and *Cdx2* is dependent on timing, responding to early (day 2-3) and not late (day 4-5) treatment with bFGF/WNT signals. The late treatment of bFGF/WNT in the D5H+ condition prevents expression of ventral neural genes *Phox2b* and *Olig2,* normally induced in the hindbrain at this timepoint (J-K, compared D5H to D5H+). By contrast, D4-5 treatment with bFGF/WNT induces dorsal and intermediate neural tube genes, indicated by *Pax7* (L) and *Dbx1* (M), respectively. (N-O) mRNA-seq expression profile of neural genes *Sox1* (N) and *Pax6* (O) indicates that neural progenitor genes are upregulated at D4, which follows treatment with neuralising signals (RA and SHH). TPM=transcripts per million.

### Chromatin accessibility defines neural progenitor identity

Genes expressed in neural progenitors can be controlled by different regulatory elements at different AP positions (Brunelli et al., 2003; Epstein et al., 1999; Uchikawa et al., 2003), therefore we hypothesized that differential enhancer usage would provide an objective definition of cell identity, and a means to follow the transition of cells into distinct neural cell types. We therefore used ATAC-seq (Buenrostro et al., 2013) to examine chromatin accessibility in cells across all stages and conditions of ESC differentiation to anterior, hindbrain or spinal cord fate (Figures 1C-D and Fig S1). As anticipated, distinct chromatin accessibility profiles were evident in different cell types. For example, enhancers directing pluripotency genes, such as *Oct3/4/Pou5f1* (Yeom et al., 1996), were accessible in ESCs but not in any of the three neural conditions, D5A, D5H or D5SC (Figure 1C). By contrast, neural enhancers, such as a previously identified enhancer of *Olig2* (Oosterveen et al., 2012; Peterson et al., 2012) showed the opposite behaviour, exhibiting accessibility in neural conditions, particularly in hindbrain and spinal cord NPs where it is highly expressed, but not in ESCs (Figure 1D). Genome wide comparisons between ESCs and neural progenitors further revealed widespread differences in accessibility between D0 ESCs and each of the D5 NPs, in line with previous observations that different tissue types present entirely different chromatin landscapes (Soucie et al., 2016; Visel et al., 2009). Compared to ESCs, D5 spinal cord NPs increased accessibility at 10,880 sites genome wide, while 13,804 sites became inaccessible in these cells. Further comparisons revealed that as cells differentiated to neural progenitors, more differences in chromatin accessibility were established (Figure 1F) and that sites open in all neural conditions (D5A, D5H and D5SC) displayed accessibility at active neural regulatory regions marked by H3K27ac (Peterson et al., 2012) (Figure 1G). Thus, ATAC-seq allows the identification of regulatory regions that define the neural progenitor lineage.

### Differences in chromatin accessibility define axial identity of neural progenitors

We next sought to identify the regulatory signatures that define neural progenitors with different AP identities. To this end, we performed unsupervised clustering using self-organizing maps (SOM) (Haberle et al., 2014; Törönen et al., 1999) of all regulatory regions that became accessible after removal of ESCs from pluripotent conditions (Figure 2A and see Methods). This allowed us to classify chromatin accessible regions that displayed different dynamics and to explore their relatedness to each other (see below). In particular, we recovered a large set of accessible regions that were common to all three neural progenitor subtypes (Figure 2A, black clusters). We mined 2.56 million DNaseI-hypersensitive sites (DHS) from the ENCODE regulatory element database (Consortium, 2012; Sloan et al., 2016), which covers regulatory sites across a range of stages and tissues. Comparison between the sites we identified in neural progenitors (Figure 2A, black clusters) and the ENCODE data demonstrated that our data were enriched with chromosomal locations associated with open chromatin in neural tissues (Figure S2A). We refer to these common sites as “neural sites”.

**Figure 2.**
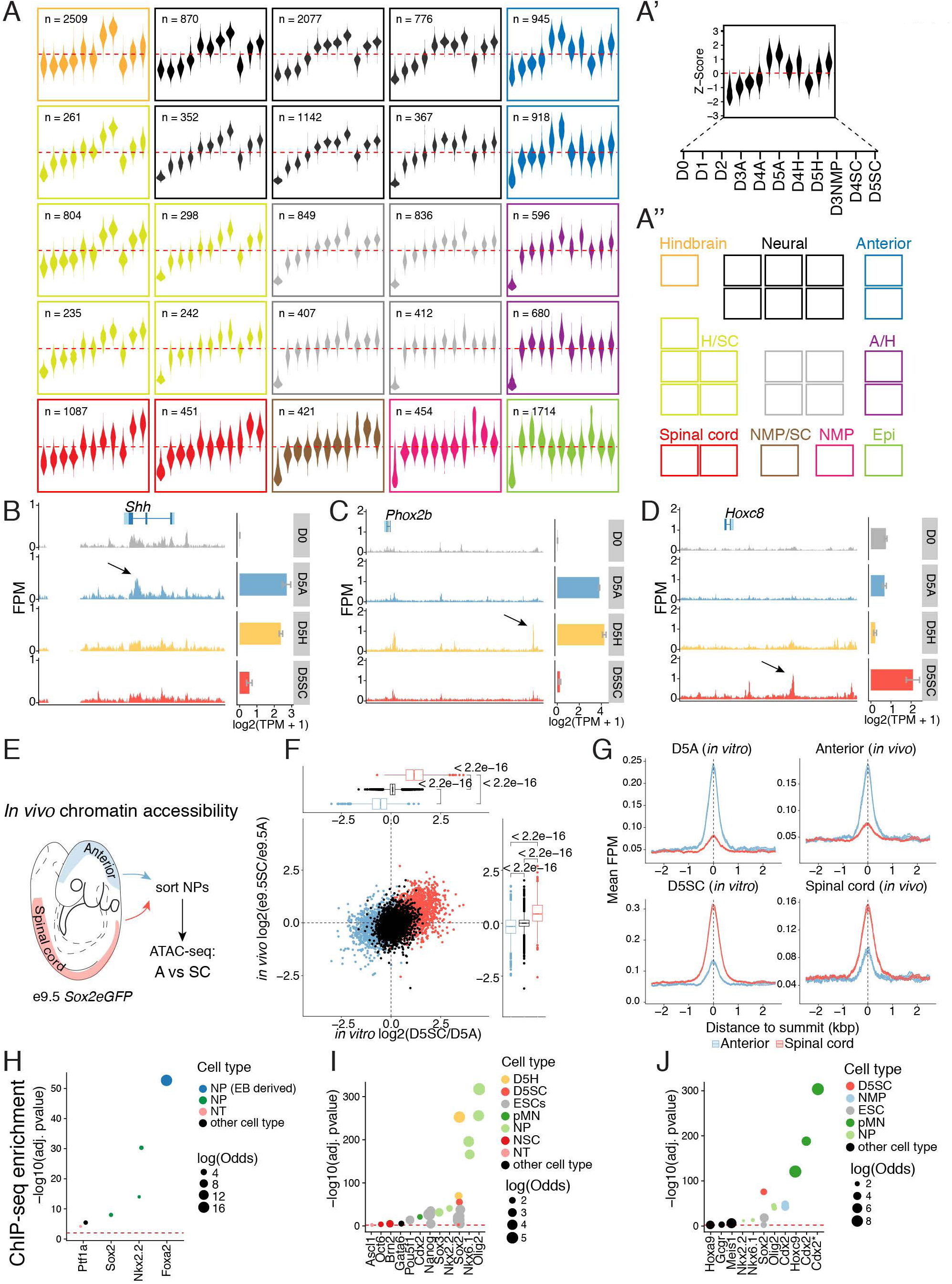
Differential enhancer usage and transcription factor engagement reveal AP identity of neural progenitors. **(A)** Self-organizing map (SOM) of all regulatory regions from all conditions and time points that show differential accessibility relative to day 0. Each plot represents the z-score for each region, across each condition (A’). Regions were classified into 10 clusters on the basis of their accessibility as outlined in A’’. Many sites are common (“neural sites”) to all neural progenitors (black cluster, n=5584), these differ from epiblast regulatory regions that are accessible at early stages of the differentiation (Epi; green cluster, n=1714). Region-specific sites are also identified in anterior (blue cluster, n=1863), hindbrain (orange cluster, n=2509) and spinal cord (red cluster, n=1538) progenitors. A distinct set of regulatory regions was observed specifically opening in D3 NMPs (pink cluster, n=454 regions). A/H represents accessible regions shared between anterior and hindbrain (lime cluster, n=1840); H/SC represents shared hindbrain and spinal cord sites (purple cluster, n=1276); NMP/SC represents shared neuromesodermal progenitor and spinal cord sites (brown cluster, n=421). Grey shaded cluster represents unclassified sites. **(B-D)** Examples of ATAC-seq accessible regions (mm10 assembly) that define anterior (B), hindbrain (C) or spinal cord (D) day 5 progenitors that were identified using the SOM presented in A and their gene expression profile determined by mRNA-seq (Error bars = SEM). Anterior progenitors display region-specific sites opening at *Shh* (B), while hindbrain progenitors demonstrate a *Phox2b* specific site (C) and spinal cord progenitors open a *Hoxc8* regulatory region (D). For a complete listing refer to Table S1. **(E-G)** *In vivo* ATAC-seq confirms a correlation between accessibility profiles found in neural cells *in vitro* and *in vivo*. E9.5 *Sox2eGFP* reporter embryos were dissected to obtain neural progenitors from anterior (blue shading) and spinal cord (red shading) regions of the neural tube (E). The fold change in accessibility at anterior (blue), and spinal cord (red) sites identified *in vitro* in spinal cord progenitors relative to anterior neural progenitors *in vivo* correlates with AP identity (F). By contrast, common neural sites identified *in vitro* (black) are similar in both populations *in vivo* and *in vitro.* Anterior sites identified *in vitro* show preferential accessibility *in vivo* in anterior compared to spinal cord progenitors (G), while spinal cord *in vitro* sites show more accessibility *in vivo* in spinal cord compared to anterior neural progenitors. (P values determined using Wilcoxon Rank Sum Test.) **(H-J)** ChIP-seq enrichment analysis of factors present in region specific sites (see Methods). Anterior regions are enriched in FOXA2 binding events (H), while hindbrain sites are enriched with OLIG2 and NKX6-1 (I). Spinal cord sites are instead enriched with CDX2 (J). SOX2 ChIP-seq in day 5 hindbrain (D5H) and spinal cord (D5SC) cells reveals that the binding site preference of this SOXB1 TF is condition-specific (I-J). Note that CDX2* denotes CDX2 ChIP-seq performed in the presence of FGF signaling (Mazzoni et al., 2013). FPM=fragments per million; neural EB=embryoid bodied-derived neural progenitors; NMP=neuromesodermal progenitors; pMN=motor neuron progenitors; NP= neural progenitors; NT=neural tube; TPM=transcripts per million.

In addition to neural sites, clustering revealed that distinct sets of regulatory regions became accessible in neural progenitors depending on their AP identity: 1863 sites were enriched specifically in anterior neural progenitors (Figure 2A, orange cluster), 2509 in hindbrain progenitors (Figure 2A, blue clusters) and 1538 in spinal cord progenitors (Figure 2A, red clusters). Examining the position of these “region-specific” regulatory sites indicated that they were also enriched for open chromatin regions enriched in neural tissues (Figure S2B-D) and displayed activity in neural tissues when compared to other tissues documented in the VISTA enhancer database (Visel et al., 2007) (Figure S2E). In contrast to common neural sites, however, which were mainly (~63.8%) located close to the transcriptional start site (TSS) of coding genes (Figure S2F), region-specific sites displayed the opposite behaviour, lying predominantly in distal regions of the genome (Figure S2G-I).

Gene-to-peak associations revealed that region-specific sites flanked many neural genes and reflected the AP identity of the cells (Figure 2B-D and Table S1). Anterior NPs displayed region-specific sites at *Shh* (Figure 2B), overlapping the previously identified Shh brain enhancer (Epstein et al., 1999). By contrast, hindbrain region-specific sites flanked the cranial MN gene *Phox2b* (Figure 2C), and in the spinal cord, region-specific sites flanked many posterior *Hox* genes expressed in the spinal cord including *Hoxc8* (Figure 2D) in addition to neural genes including *Nkx6-1, Lhx1, Sox11* and *Sox2* (Table S1 lists all peak coordinates for region-specific sites and associated genes).

To test whether the different region-specific signatures observed *in vitro* reflect differences present *in vivo*, we performed ATAC-seq on mouse NPs isolated from different AP levels: anterior (forebrain/midbrain/anterior hindbrain levels) or spinal cord levels (cervical/thoracic) of e9.5 mouse embryos (Figure 2E-G). This developmental time point corresponds most closely to the gene expression profile of D4-5 NPs (Gouti et al., 2017). To isolate neural progenitors from the surrounding tissue, we used *Sox2eGFP* reporter mice that express GFP throughout the nervous system (Ellis et al., 2004) to obtain distinct anterior and posterior regions of the neural tube (Figure 2E and see Methods). To ask if AP differences observed *in vitro* predict AP identity *in vivo*, we examined the change in accessibility of *in vitro*-defined spinal cord sites, relative to in vitro-defined anterior sites, under *in vivo* conditions (Figure 2F). Strikingly, region-specific signatures showed an enrichment that reflected their AP identity *in vivo* similar to that observed *in vitro*, whereas neural sites common to all *in vitro* derived neural progenitors were equally enriched in both populations *in vivo* (Figure 2F). Anterior *in vivo* NPs demonstrated increased accessibility at sites that define *in vitro* anterior neural identity and not spinal cord (Figure 2G). For example, regulatory regions included sites that flanked neural genes, such as *Shh,* where accessibility overlapped the *Shh* brain enhancer (Epstein et al., 1999), under anterior but not spinal cord *in vivo* conditions (Figure S2J, arrow). Similar results were obtained when examining *in* vivo-derived spinal cord NPs: increased accessibility was observed in regions that define spinal cord NPs *in vitro* (Figure 2G). For example, accessibility was observed at the *Olig2* enhancer (Oosterveen et al., 2012; Peterson et al., 2012), which is repressed in anterior neural conditions (Figure S2K, arrow). In summary, changes in chromatin accessibility reflect the AP identity of neural progenitors of the nervous system and are recapitulated by NPs generated *in vitro*.

### Context-dependent binding of neural TFs defines axial identity

We next interrogated the *in vitro* region-specific signatures of anterior, hindbrain and spinal cord progenitors to identify TFs that recognise these AP-specific sites. We performed ChIP-seq enrichment analysis using a set of publicly available datasets (Sheffield and Bock, 2015) covering 910,490 regulatory regions from 270 different TFs (Table S2). Our analysis revealed that distinct TF binding was enriched in the three distinct subtypes: anterior NPs were enriched with FOXA2 and NKX2-2 binding sites (Figure 2H); by contrast, hindbrain sites contained OLIG2 and NKX6-1 (Figure 2I); and spinal cord sites were enriched for CDX2 and HOXC9 binding (Figure 2J). Motif enrichment with Homer (Heinz et al., 2010) predicted an enrichment of SoxB1 TF motifs (SOX1/2/3), in all three neural progenitors subtypes (Figure S3A-C), consistent with their expression throughout the neuraxis and central role in neural development (Avilion et al., 2003; Kamachi and Kondoh, 2013; Wood and Episkopou, 1999; Ying et al., 2003). Notably, hindbrain and spinal cord cells, which are both exposed to the same signals (RA/SHH) from D3-D5 are enriched for SOXB1 binding but at distinct genomic sites (Figure S3B-C). The presence of posterior HOX binding events together with SOX in spinal cord progenitors suggested that region-specific expression of *Hox* genes influenced the binding site preference of the core neural SOXB1 TFs. Likewise, posterior *Hox* genes can alter the binding site preference of SOX factors when misexpressed in the cortex (Hagey et al., 2016).

To validate these in-silico findings, we performed ChIP-seq of the SOXB1 TF, SOX2, in D5 hindbrain and spinal cord NPs (Figure 2I-J). This confirmed hindbrain predicted SOX sites were physically engaged with SOX2 in hindbrain NPs (Figure 2I) and not spinal cord NPs at day 5 of the differentiation (Figure 2J). Conversely, SOX2 accessible sites specific to the spinal cord condition showed increased engagement of SOX2 in spinal cord versus hindbrain conditions (Fig.2I-J). Collectively, these data demonstrate the utility of ATAC-seq for predicting factors enriched at region-specific sites in NPs. Furthermore, it demonstrates that NPs develop region-specific transcription factor binding at a time when they are exposed to the same extrinsic signals - RA/SHH. This raises the question of how differences in regional identity and transcription factor engagement are established.

### Posteriorising signals that promote spinal cord identity depend on developmental timing

We took advantage of the temporal resolution afforded by the *in vitro* differentiation to define when AP identity is established in neural progenitors. To address this question, we examined when region-specific regulatory regions became accessible, and under which conditions this occurred (Figure 3A-C). The prevailing view is that to generate neural cells with a spinal identity, anterior neural cells must be gradually posteriorised to acquire a spinal fate (Davis-Dusenbery et al., 2014; Stern, 2001). To test this assumption, we asked if spinal cord progenitors transitioned via an anterior or hindbrain identity, before acquiring spinal cord identity. We assessed region-specific sites and their behaviour over the course of the differentiation. In contrast to previous models, we found that spinal cord cells failed to exhibit transient accessibility at either anterior (Figure 3A) or hindbrain (Figure 3B) region-specific sites, challenging this view. Instead, spinal cord-specific sites became accessible in spinal cord conditions by D4 of the differentiation (Figure 3C).

Examining the timing of region-specific sites further revealed a synchronicity between neural progenitor identity and the establishment of AP fate in cells. Specifically, anterior, hindbrain and spinal cord progenitors begin to exhibit region-specific accessibility between D3-4, coincident with the emergence of neural sites (Figure 2A, black cluster). Both the hindbrain and spinal cord progenitors are exposed to the same conditions (RA + SHH; Figure 1A) yet adopt region-specific signatures at this same time point (Figure 3B-C). Thus, extrinsic signals present at the time the regional signatures emerge are not sufficient to promote distinct regulatory element usage in cells. The only difference in generating hindbrain vs spinal cord progenitors *in vitro*, is the addition of WNT signals that is added together with FGF between D2-3 of the spinal cord differentiation (Figures 1A and 3D). This suggests that WNT signalling, together with FGF, plays a critical role in posteriorising cells to adopt spinal rather than hindbrain fates, consistent with previous studies (Gouti et al., 2014; Nordström et al., 2006). Our data suggest that at the genomic level, this requires the suppression of hindbrain sites in response to RA/SHH signals, and adoption of a distinct set of SC specific sites (Figures 2A and Figure 3B-C).

While the specific timing and onset of neural progenitor fates has been difficult to define *in vivo*, we took advantage of the in vitro system to alter the timing of extrinsic signals to directly test the temporal requirement for WNT signalling. To this end, we provided the same 24h pulse of FGF/WNT but at later stages of the differentiation, to hindbrain progenitors at D4-5 (Figure 3D’). We then asked whether exposure to these signals at this time was sufficient to promote a spinal cord instead of hindbrain identity (Gouti et al., 2014; Nordström et al., 2006). Altering the timing of FGF/WNT from D2-3 to D4-5 (Figure 3D’) resulted in the induction of canonical WNT signalling target genes such as *Notum*, to levels comparable with the induction observed at D3 when provided to cells between D2-3 (Figure 3E, compare D3NMP and D5H+). However, shifting the treatment of FGF/WNT to D4-5 was no longer sufficient to induce expression of posterior *Hox* genes characteristic of the spinal cord condition, including *Hoxc8, Hoxb9*, which are induced from D3 following D2-3 FGF/WNT treatment (Figure 3F-G). Likewise, we found that the induction of *Brachyury* (*T/Bra*) and *Cdx2*, normally induced at D3 by a D2-3 pulse of FGF/WNT (Gouti et al., 2014) (Figures 3H-I and S4A-C), was no longer observed at D5 following FGF/WNT treatment between D4-5 (Figure 3H-I, compare D3NMP and D5H+).

The failure of hindbrain progenitors to upregulate spinal cord genes suggests that administration of FGF/WNT signals at this stage is not capable of posteriorising cells. We further investigated the neural identity of the cells resulting from FGF/WNT treatment between D4-5. We found that the expression of the ventral neural markers *Phox2b* and *Olig2*, normally expressed in hindbrain conditions (Figure 1B), was no longer maintained following FGF/WNT treatment in these cells (Figure 3J-K). By contrast, expression of *Pax7* and *Dbx1*, markers of dorsal and intermediate neural tube fates, respectively, was observed (Figure 3L-M). WNT signalling is known to promote dorsal neural fate at the expense of ventral fate in the neural tube (Lei et al., 2006; Wang et al., 2011). Thus, the later treatment of cells with FGF/WNT, a point at which cells have begun expressing neural progenitor markers such as *Sox1* and *Pax6* (Fig. 3N,O) in response to neuralising signals provided from D3 (Figure 1A), is consistent with WNT promoting dorsal (*Pax7, Dbx1*) at the expense of ventral (*Phox2b, Olig2*) neural cell fates during neural tube development (Alvarez-Medina et al., 2008; Lei et al., 2006; Muroyama et al., 2002; Wang et al., 2011). The posteriorising activity of WNT together with FGF signalling is thus dependent on precise developmental timing: prior to neural progenitor establishment (D2-3), these signals are capable of posteriorising cells, yet after cells commit to the neural lineage (D4-5), the same combination of extrinsic signals promotes dorsal in place of posterior cell fates in the nervous system (Le Dréau and Martí, 2012; Gouti et al., 2015) Taken together these data indicate that the generation of spinal cord fates does not follow a gradual posteriorisation of more anterior neural progenitors, as cells lose the competency to be posteriorised following establishment of neural fate.

### WNT directs a transient set of chromatin remodelling events

To understand how FGF/WNT signals exert a stage-specific, posteriorising effect in cells prior to neural progenitor establishment, we examined the chromatin landscape in cells following FGF or FGF/WNT treatment at D3. We found that at D3, cells that had been exposed to FGF/WNT for 24h displayed accessibility at 875 unique regions (Figure 2A, NMP/SC and NMP cluster). Strikingly, of these 875 sites, 454 (51.8%) were immediately downregulated as cells committed to spinal cord fates by D5 (Figure 2A, NMP cluster). Thus, as cells transition to a spinal cord identity, they transiently adopt a genomic signature, in response to FGF/WNT signals, that is distinct from both the epiblast (Figure 2A, epiblast cluster) and neural regulatory signatures (Figure 2A, black clusters). These cells include the bipotential population of NMPs, which contributes to both the spinal cord and somites (Gouti et al., 2014; Tsakiridis et al., 2014; Turner et al., 2014; Tzouanacou et al., 2009; Wymeersch et al., 2016), that expresses the transcription factors Sox2, *T/Bra* and *Cdx1,2,4* (Martin and Kimelman, 2012; Olivera-Martinez et al., 2012; Tsakiridis et al., 2014; Young et al., 2009).

To further validate the *in vitro* signature, we asked to what extent these sites overlap with the *in vivo* chromatin accessibility associated with NMPs. We took advantage of existing ATAC-seq data collected from whole mouse epiblasts from E6.0-7.2 and from E7.5 posterior mouse tissue – the tissue which contains NMPs (Neijts et al., 2016). We found that more than 71% of sites induced by FGF/ WNT signalling *in vitro* at D3 overlapped with accessible sites found in E7.5 posterior mouse embryos (Figure 4A), while the overlap with either the epiblast (29% at E6.0) or purified NPs from E9.5 embryos is much less (less than 5%, this study; Figure 4A). We also found that the epiblast-specific sites defined in the self-organizing map (Figure 2A, epiblast cluster) showed the greatest overlap with those found *in vivo* in E6.0 epiblast (Figure 2A), while very little overlap was evident with *in vivo* neural progenitors (Figure 4A). These data suggest that the epiblast and NMP signatures identified *in vitro* correspond to their respective, tissue-specific, regulatory signatures *in vivo*.

**Figure 4.**
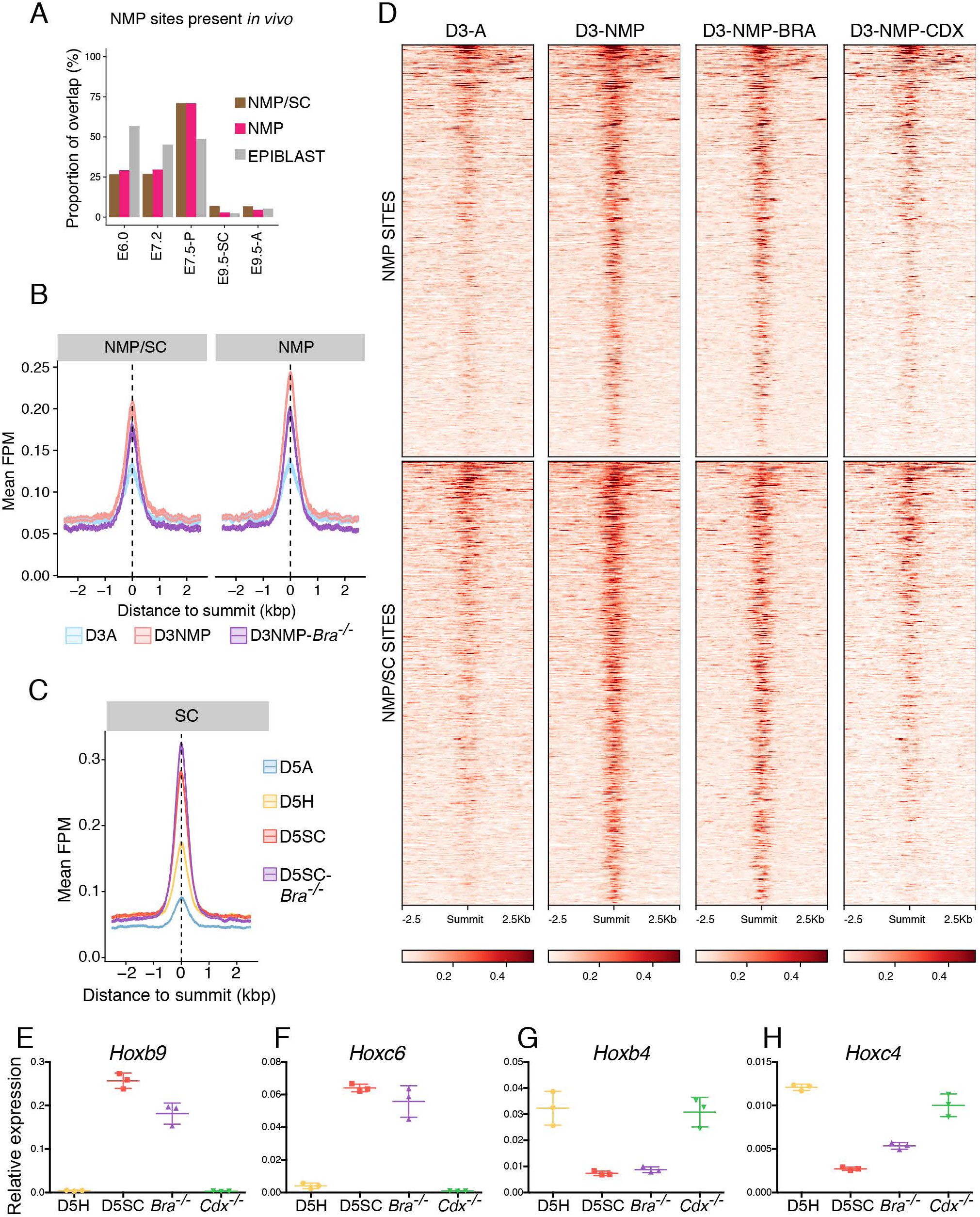
*T/Bra* is dispensable for the chromatin remodeling events that establish NMP and SC identity. **(A)** The proportion of NMP, NMP/SC shared or epiblast genomic sites identified *in vitro* and their overlap with accessible regions present *in vivo*. The highest proportion (71%) of NMP site overlap coincides with the posterior E7.5 embryo (E7.5-P) which harbours NMPs (Neijts et al., 2016) and this contrasts with neural progenitors which lack this signature in the spinal cord (E9.5-SC) and anterior nervous system (E9.5-A; this study). (**B**) The average accessibility profile of NMP/SC shared and NMP specific sites in wild-type and *T/Bra^-/-^* cells. Accessibility of these sites remains largely unchanged in cells lacking *T/Bra^-/-^* cells at day 3 of the spinal cord differentiation. (**C**) *T/Bra^-/-^* mutant cells differentiated to D5 under spinal cord conditions retain accessibility at spinal cord (SC) genomic sites. (**D**) Heatmap showing the accessibility of NMP and NMP/SC (neuromesodermal progenitors and spinal cord) shared sites is maintained in the absence of *T/Bra* (*Bra^-/-^*) but is dramatically reduced in the absence of all three *Cdx* TFs, *Cdx1,2,4* (*Cdx^-/-^*). (**E-H**) RT-qPCR of *Hox* genes at D5 of the *in vitro* differentiation shows the difference in AP identity between hindbrain (D5H) and spinal cord (D5SC) cells under wildtype conditions, compared with *T/Bra^-/-^*mutant (*Bra^-/-^*) and *Cdx^1,2,4-/-^* mutant (*Cdx^-/-^*) cells differentiated under spinal cord conditions. Note that *Bra^-/-^* mutant cells retain expression of spinal cord *Hox* genes *Hoxb9* (E) and *Hoxc6* (F) in contrast to *Cdx* mutant cells which fail to upregulate these genes and instead express *Hoxb4* (G) and *Hoxc4* (H), which occurs in hindbrain conditions. FPM= fragments per million; NMP= neuromesodermal progenitors; pMN= motor neuron progenitors.

As NMPs generate paraxial mesoderm and are present in posterior tailbud tissues, it remained a possibility that the NMP sites we had recovered *in vitro* represented a nascent mesodermal population. Both NMP and paraxial mesodermal progenitors express *T/Bra* (Garriock et al., 2015 Tsakiridis et al., 2014; Wymeersch et al., 2016). Therefore, we examined chromatin accessibility in ESCs in which *T/Bra* had been genetically inactivated. ESCs lacking *T/Bra* are able to generate spinal cord progenitors, but are incapable of forming paraxial mesoderm (Gouti et al., 2014). Strikingly, the loss of *T/Bra* had a negligible effect on both the NMP/SC shared and NMP-specific signature induced by WNT exposure (Figure 4B), as NMP sites remained accessible in mutant cells (Fig. 4B,D). Furthermore, T/Bra-lacking ESCs differentiated to D5SC maintained spinal cord chromatin accessible sites (Figure 4C) and maintained the expression of posterior *Hox* genes such as *Hoxb9* and *Hoxc6* (Figure 4E-F), while expression of *3’ Hox* genes (like *Hoxb4* and *Hoxc4)* was reduced, similar to WT D5SC cells (Figure 4G-H). Thus, T/BRA, which is necessary for paraxial mesoderm specification (Nowotschin et al., 2012; Rashbass et al., 1991), is not responsible for the chromatin remodelling events associated with NMP identity (Figure 4B,D), nor is it required to induce posterior *Hox* genes or spinal cord identity (Figure 4C, E-F) (Gouti et al., 2014).

### The acquisition of spinal cord fate requires CDX to repress hindbrain identity

To establish which factors are responsible for mediating WNT-dependent chromatin remodelling events, we performed ChIP-seq enrichment analysis on NMP-specific sites (Figure 5A). In this analysis, we found that the NMP regulatory regions are highly enriched with CDX2 TF binding events (Figure 5A). Nucleotide resolution analysis of the frequency of transposon-mediated integration events further verified the presence of a CDX “footprint” (Buenrostro et al., 2013) present in the chromatin landscape of D3NMP cells. This suggested physical engagement of these factors occurs at sites of open chromatin (Figure S4D) and supports the idea that CDX plays an important role downstream of WNT signalling to promote NMP identity. Recent observations support these findings, suggesting that CDX factors promote a niche that sustains growth of the posterior tailbud and *in vitro*-derived NMPs (Amin et al., 2016). However, the genetic removal of *Cdx* TFs or combined absence of *Cdx2* and *T/Bra* results in severe axis elongation *in vivo* (Amin et al., 2016; van Rooijen et al., 2012a; Young et al., 2009). This has precluded a direct analysis of the cellular context required for these factors to function *in vivo*, and hence their role during spinal cord generation has remained unclear.

**Figure 5.**
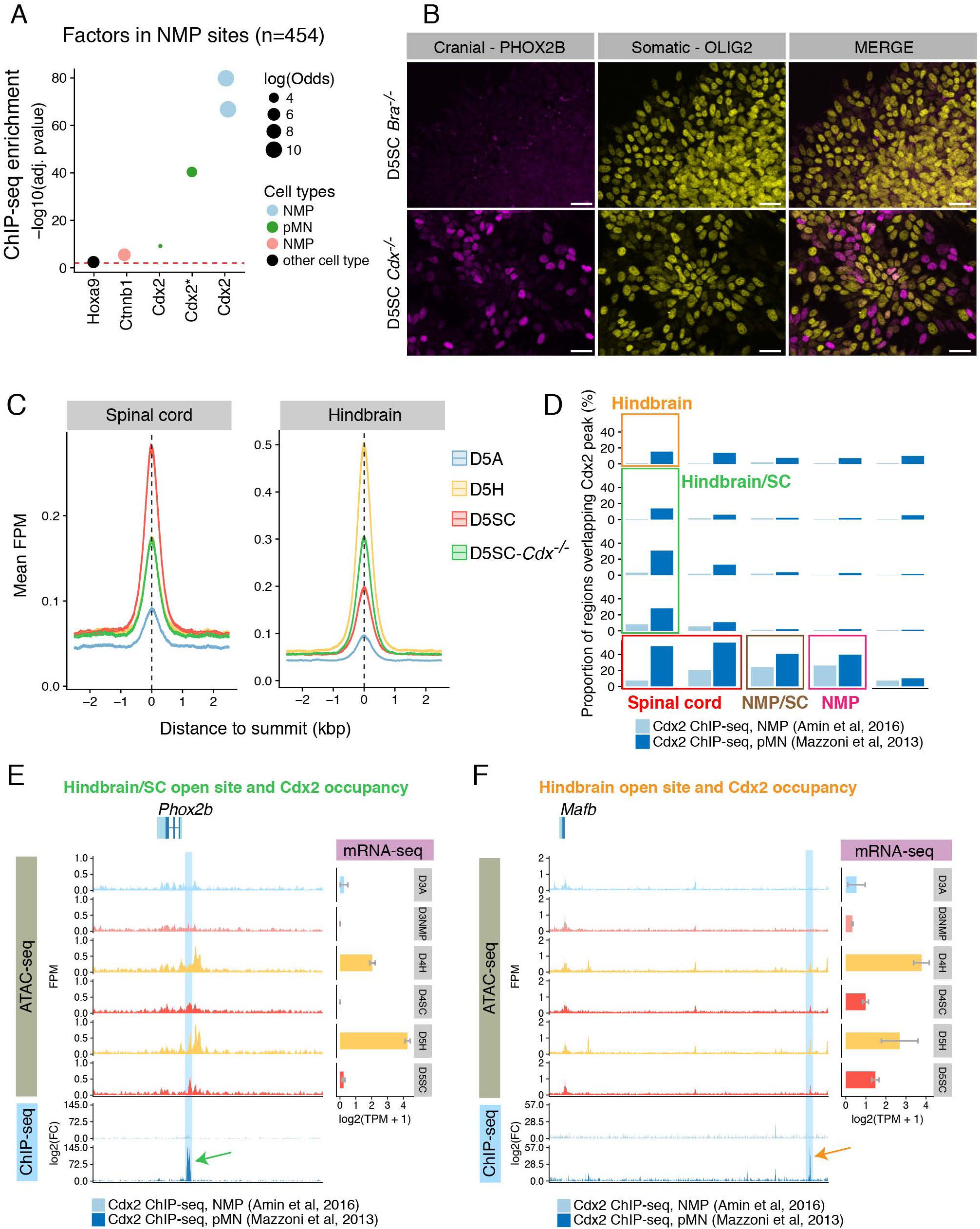
Cdx transcription factors remodel the chromatin landscape to posteriorise cells and repress cranial motor neuron identity. **(A)** ChIP-seq enrichment analysis reveals that CDX2 is highly enriched at NMP specific sites. (**B**) Removal of the three *Cdx* transcription factors *Cdx1/2/4* results in ectopic production of cranial motor neuron progenitors marked by PHOX2B, that are normally repressed in spinal cord conditions. Scale bars = 20 microns. (**C**) The average profile of spinal cord specific sites (left plot) shows that relative to day 5 spinal cord (D5SC, red), these sites are lost in *Cdx* mutant cells differentiated under the same conditions (D5SC-Cdx^-/-^, green), to the same extent observed in D5 hindbrain cells (yellow). Under spinal cord conditions, *Cdx* mutant cells show increased accessibility at hindbrain specific sites (right plot). (**D**) The proportion of accessible regions bound by CDX2, as indicated by CDX2 ChIP-seq analysis from neuromesodermal progenitors (NMP, Amin et al., 2016) and motor neuron progenitors (pMNs, Mazzoni et al., 2016) derived *in vitro*, demonstrates an enrichment for CDX2 at NMP, NMP/SC (NMP and spinal cord shared) and spinal cord (SC) sites. Furthermore, under pMN conditions, CDX2 targets accessible regions that are shared between the hindbrain and spinal cord. (**E**) At the *Phox2b* genomic region, a hindbrain/spinal cord shared site is bound by CDX2 in pMN conditions (blue shading). (**F**) A hindbrain-accessible site is bound by CDX2 (blue shading) at the *Mafb* locus in pMN conditions.

We took advantage of the *in vitro* differentiation, where we could directly test the function of all three CDX TFs in both the generation of NMPs and spinal cord cell types. Using an ESC line lacking *Cdx1,2,4* (*Cdx^1,2,4-/-^*) (Gouti et al., 2017), we asked if WNT treatment was sufficient to remodel the chromatin landscape, as observed in NMPs lacking *Bra* (Figure 4B). In contrast to the loss of *T/Bra,* however, the elimination of all three *Cdx* TFs had a profound effect on the response to WNT signalling (Figure 4D). Only a small fraction of the NMP and NMP/SC shared sites remained accessible in *Cdx^1,2,4-/-^* cells with most NMP sites displaying similar accessibility to D3A cells that were not treated with WNT signals (Figure 4D). Similarly, the CDX footprint observed in WT cells was no longer observed (Figure S4D). This suggests that CDX factors are essential for the remodelling of chromatin accessibility associated with an NMP state (Amin et al., 2016) as well as the transition from an NMP to spinal cord fate (Figure 4D). Furthermore, the differentiation of mutant *Cdx* cells to neural progenitors no longer resulted in the expression of spinal cord *5’ Hox* genes *Hoxb9, Hoxc6* (Figure 4E-F). Instead, *Cdx* mutant cells expressed *Hoxb4, Hoxc4*, as observed in WT hindbrain conditions at D5 (Fig. 4G-H).

These findings suggest that in the absence of *Cdx* TFs, the application of posteriorising signals no longer promotes a posterior neural identity in cells, and thus the generation of more anterior neural fate ensues. To test this prediction, we performed immunofluorescence on *Cdx^1,2,4-/-^* mutant cells at D5 of the spinal cord differentiation, to confirm the complement of MN subtypes present at this time. We found ectopic generation of visceral MNs marked by PHOX2B in *Cdx* mutant cells (Figure 5B), which normally occurs in hindbrain but not spinal cord conditions (Figure 1B). In addition, sustained *Olig2* induction was observed suggesting the removal of CDX TFs does not impede neural progenitor establishment, per se (Figure 5B). In agreement with this observation, analysis of the chromatin accessibility landscape present in D5 *Cdx^1,2,4-/-^* cells differentiated under spinal cord conditions revealed that these cells lacked spinal cord identity sites and instead, gained accessibility at hindbrain identity sites (Figure 5C). These data demonstrate that CDX factors are required to suppress the accessibility of hindbrain identity sites in response to RA/SHH signals, and allow the specific acquisition of spinal cord accessible regions.

To address how CDX factors are capable of both repressing hindbrain accessible sites and allowing the induction of a spinal cord regulatory program, we took advantage of previously published CDX2 ChIP-seq from *in vitro*-generated NMPs (Amin et al., 2016) and motor neuron progenitors (pMNs) (Mazzoni et al., 2013) and mapped the proportion of bound sites in the accessible regions clustered in the self-organising map (Figure 2A) of ATAC-seq accessible regions (Figure 5D). This allowed us to define direct targets of CDX2 (Table S3) and monitor in which cell types these regions were accessible. As expected, CDX2 binding was highly enriched at genomic sites accessible in NMPs, as well as SC specific sites and sites found in both conditions (Figures 5D and S5A). This overlap suggests a critical role for CDX factors in the transition to a spinal neural identity, where it targets many regions of the posterior *Hox* genes (Table S3) (Amin et al., 2016; Mazzoni et al., 2013; Neijts et al., 2017; Young et al., 2009). More strikingly, however, we found that CDX2 bound to additional target sites, outside of the *Hox* locus, which were accessible in hindbrain and spinal cord conditions (Figures 5D, green clusters and S5A), as well as sites exclusive to the hindbrain (Figures 5D, orange cluster and S5A). Examination of these CDX2-bound sites revealed a shared hindbrain/spinal cord site lying upstream of *Phox2b* (Figure 5E and TABLE S3 list of regions). This demonstrates that CDX factors are capable of directly targeting neural genes, including those that are repressed from spinal cord conditions. Similarly, previous studies have suggested that CDX1 directly binds to regulatory elements at *Mafb*, and represses the expression of this hindbrain marker (Kim et al., 2005; Sturgeon et al., 2011). Consistent with this we found that CDX2 directly targets *Mafb* at a region only accessible under hindbrain conditions (Figure 5F). A systematic analysis further revealed that a substantial number of hindbrain genes are repressed by CDX, as demonstrated by mRNA-seq analysis *in vivo* (Figure S4E) (Amin et al., 2016). This includes *Mafb* and the binding of CDX2 correlates with preventing accessibility at a nearby regulatory region, as *Cdx* mutant cells show striking increases in accessibility at this site from D4-5 under spinal cord conditions (Figure S4F, arrows). Gene ontology analysis of the major pathways and processes enriched at CDX2-engaged sites (Figure S5B-E) confirmed that CDX2 occupancy at hindbrain-accessible sites is directly linked with neural genes (Figure S5C), whereas in both NMP/SC and SC the CDX2 bound accessible sites were associated with genes implicated in anteriorposterior patterning (Figure S5D-E).

In summary, genome-wide analyses indicate that in addition to establishing NMP identity and driving activation of posterior *Hox* genes (Amin et al., 2016; Mazzoni et al., 2013; Neijts et al., 2017), CDX factors play a central role in directly targeting neural genes at regulatory sites associated with hindbrain identity (Figure 5D-F). We propose that this mechanism ensures the repression of hindbrain genes in response to RA/SHH signals, in addition to the priming of posterior *Hox* genes, which drive spinal cord identity. The induction of CDX, prior to neural induction, is therefore critical in establishing the appropriate SC-specific regulatory signature, while further repressing hindbrain fate in response to neuralising signals. We propose that this dual functionality of CDX is essential to restrict the generation of specific neuronal subtypes to discrete AP levels of the neural tube. Such a mechanism ensures the production of, for example, visceral MNs in the posterior hindbrain but not the spinal cord (Figure 6A).

**Figure 6.**
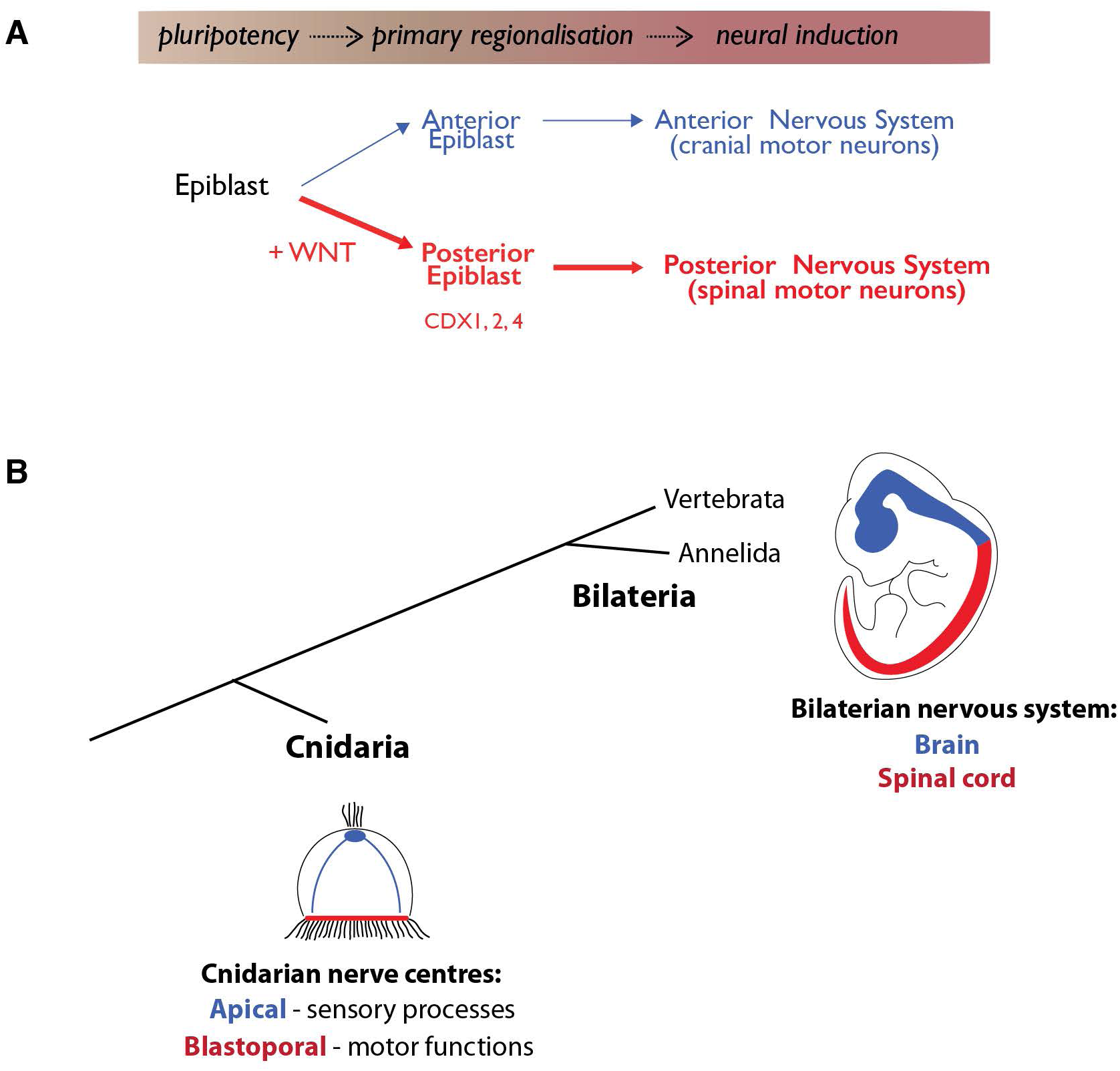
Proposed model of nervous system development. **(A)** Pluripotent epiblast cells in the early embryo are first allocated into anterior (blue) or posterior (red) populations before they have committed to a neural identity. When cells undergo neural induction to form the progenitors of the nervous system, these two populations give rise to distinct neural subtypes. Cells which have been posteriorised form spinal neural cell types (e.g. somatic motor neurons) that make up the posterior nervous system. By contrast, anterior epiblast cells can generate cranial motor neurons, and thus, unlike posterior epiblast cells, support the generation of the anterior nervous system. **(B)** Comparisons between Cnidarian and Bilaterian animals provides supporting evidence for the dual evolutionary origin of the vertebrate central nervous system which is proposed to have arisen in a pre-bilaterian animal ancestor (Arendt et al., 2016). Cnidarians display two distinct nerve centres: apical (blue) and blastoporal (red). Blastoporal centres show expression of putative CDX orthologues (Arendt et al., 2016; Ryan et al., 2007). In Bilaterians, these separate nerve centres have expanded and merged. We speculate that the transition between these two nervous systems lies at the level of the posterior hindbrain/anterior spinal cord.

## DISCUSSION

The prevailing view of nervous system formation, summarised in the “activation-transformation hypothesis”, proposes that nervous system induction occurs in two phases (Eyal-Giladi, 1954; Nieuwkoop, 1952; Nieuwkoop and Nigtevecht, 1954; Toivonen and Saxen, 1968). In the first step, ectoderm is induced to become anterior neural tissue. Following this “activation” step, posteriorising signals convert – “transform” – anterior neurectoderm into the complete range of positional identities that comprise the neuraxis (Stern, 2001). In this view of neural induction, the generation of posterior regions of the nervous system, such as the spinal cord, require that cells first acquire forebrain identity before being caudalised to a posterior fate. Despite the widespread acceptance of this model, previous studies have lacked the cellular and temporal resolution necessary to test this model. To address this, we took advantage of *in vitro* differentiation of ESCs to assay the chromatin landscape as cells transition from pluripotency to different axial levels of the nervous system. This revealed that cells destined to form spinal cord progenitors do not transiently adopt an anterior neural state or acquire their regional identity via a gradual caudalisation of more rostral cell types (Figure 3A-C). Instead, spinal cord cells are regionally restricted prior to their commitment to a neural fate. We provide evidence that CDX remodels the chromatin landscape in cells before neural fate is established. This step is essential for the specification of a spinal cord identity in cells, and the repression of cranial MN fates (Figure 5C). Thus, specification of spinal cord fate involves cells committing to an axial identity (Figure 6A, “*primary regionalisation*’’) prior to neural induction, reversing the sequence of events implied by the ‘activation-transformation’ hypothesis and prompting a revision in our understanding of nervous system regionalization (Figure 6A).

### Regulatory signature dynamics argue against “activation-transformation”

Support for the activation-transformation hypothesis originated in embryological and molecular experiments in chick and frog embryos. For example, explants of posterior axial tissue promote midbrain and hindbrain fates from prospective forebrain tissue (Cox and Hemmati-Brivanlou, 1995) and manipulating WNT, FGF and/or RA signalling in neural plate explants alters rostral caudal identity of neural cells in ways consistent with a graded caudalising activity (Kolm et al., 1997; Lamb and Harland, 1995; Muhr et al., 1999; Nordström et al., 2006; Wilson et al., 2001). It is notable that in many of these studies the most caudal markers assayed were representative of the hindbrain or anterior spinal cord and the results were subsequently extrapolated to apply to the entire length of the spinal cord without explicit testing. While RA exposure to neural progenitors is sufficient to posteriorize anterior neural cells to form hindbrain, the most caudal identity generated in these assays corresponds to cervical (anterior) spinal cord (Gouti et al., 2014; Liu et al., 2001; Mahony et al., 2011; Maury et al., 2014; Mazzoni et al., 2013; Niederreither et al., 2000). Furthermore, treatment of anterior neural progenitors with increasing concentrations of WNT fails to caudalise these cells to a spinal cord fate, instead their identity corresponds to the posterior hindbrain (Kirkeby et al., 2012). Thus, the activation-transformation hypothesis seems compatible with the experimental evidence for regionalisation of the fore-, mid- and hindbrain, but extending the model to the spinal cord does not appear to be supported by the data (Lamb and Harland, 1995).

To systematically define neural cell identity, with the resolution to distinguish AP-specific differences, we took advantage of ATAC-seq (Buenrostro et al., 2013, 2015). We reasoned that enhancer usage, read out by chromatin accessibility, would provide an unbiased means to follow the induction of neural fate and to determine when specific regional identities are established in cells. Using a data-driven approach, we clustered regulatory regions based on their pattern of accessibility over developmental time during the formation of anterior, hindbrain and spinal cord progenitors. This revealed that a common set of regulatory regions becomes accessible across all neural progenitors. In addition, many of these sites overlap with accessible regions present in neural tissues. Thus, a distinct regulatory signature defines the neural lineage. Overlaid with this, we find that neural progenitors display additional regulatory signatures that define their AP identity, even in hindbrain and spinal cord progenitors, where their AP signatures are laid down while exposed to the same neuralising conditions (Figure 2A). Comparisons between *in vitro* and *in vivo*-derived neural cells demonstrated that the regulatory signatures observed *in vitro* reliably predicted AP position within the neural tube (Figure 2E-G). Taking advantage of the temporal resolution afforded by the *in vitro* differentiation, we traced the emergence of genomic signatures during neural induction (Figure 3A-C). This revealed that AP-specific regulatory regions appear in cells concomitant with the establishment of neural identity (from Day 3-5). Crucially, the pattern of accessibility that defined spinal cord identity (Figure 3C) appeared at the same time as the broader neural signature, as shown by the neural set of regions that open from Day 4-5 across all progenitor subtypes (Figure 2A, black clusters show neural sites). This coincides with the addition of RA and SHH – signals that promoted neural induction (Figure 3N,O). In addition, cells differentiating to a spinal cord fate did not display a transient anterior or hindbrain regulatory landscape prior to the induction of the spinal cord signature (Figure 3C). These data argue against the idea that the spinal cord is generated by posteriorising more anterior neural cells and therefore, alternative mechanisms must be involved.

### AP identity is established before the acquisition of neural identity

*In vitro*, we found that the regulatory signatures that define different regional NPs is established at the same time. Moreover, different AP identities emerge in cells under the same conditions: hindbrain and spinal cord signatures appear in cells during the period in which they are exposed to RA & SHH (Figure 3B-C). Hence, the distinction between regional identities is established before this time, when spinal cord (but not hindbrain) fated cells receive FGF/WNT signalling (Figure 1A). We tested the importance of this timing by altering when the signals were applied during neural differentiation (Figure 3D,D’). This indicated that delaying addition of FGF/WNT signals until after neural identity had been established was unable to convert hindbrain cells to a spinal cord identity (Figure 3F-G). Thus, cells must receive these signals before neural fate is established, as the competence of cells to form spinal cord is lost following neural induction. This suggests that FGF/WNT signalling establishes a posterior genomic programme in cells – a ‘primary regionalisation’ – before neural induction.

To determine what factors might determine posterior competency in cells, we examined the chromatin landscape and identified regions that responded to FGF/WNT at D3 (NMP sites; Figure 2A). These NMP sites were enriched in CDX2 TF binding sites (Figure 5A) and we identified the presence of a CDX footprint (Figure S4D) suggesting CDX factors physically engage with open chromatin sites following FGF/WNT treatment. Our findings are consistent with the established role of CDX in promoting the formation of posterior embryonic development, downstream of FGF and WNT signalling (Amin et al., 2016; Skromne et al., 2007; Young et al., 2009). The absence of CDX factors *in vivo* severely truncates embryo elongation resulting in a lack of trunk tissue (Amin et al., 2016; Chawengsaksophak et al., 2004; Young et al., 2009). It has therefore proved difficult to define the cellular context required for CDX to function, and to address how CDX may regulate the production of spinal cell types. Taking advantage of the *in vitro* system, we showed by mutating all three *Cdx* genes that these factors are necessary for remodelling chromatin and the presence of a CDX footprint (Figure S4D), making available 875 sites genome wide in response to FGF/WNT signaling (Figures 2A and 4D). Furthermore, we uncovered that the majority of these NMP sites overlap with *in vivo* accessible regions in the posterior mouse tailbud, the tissue in which NMPs reside (Figure 4A). A similar role for CDX was recently proposed in NMPs generated *in vitro* from mouse EpiSCs (Amin et al., 2016; Neijts et al., 2016). Our data agree and extend these findings by demonstrating that cells lacking *Cdx1/2/4* can no longer generate a genomic signature associated with NMP identity (Figure 4D) and ultimately generate hindbrain NPs instead of spinal cord (Figure 5C-D).

CDX factors are crucial to prevent accessibility at hindbrain specific sites in addition to the maintenance of NMP identity. Loss of *Cdx* function has been shown to promote hindbrain fate at the expense of spinal cord in zebrafish (Skromne et al., 2007) and in mouse (Young et al., 2009). In the latter case, *Hox* genes are mostly capable of recovering the posterior elongation defects observed in these mutants. However, in zebrafish, overexpression of posterior *Hox* genes was incapable of rescuing the defect under these conditions. It has therefore remained difficult to determine the precise function of CDX proteins, the timing and cell type in which they function, and how they exert their effect *in vivo*. We resolve this uncertainty and show that a limited developmental window exists, between pluripotent epiblast cells and neural progenitors, in which FGF/WNT can induce *Cdx* expression (Figures 3I and S4A-C) and thus exert posteriorising effects (Figure 3F,G). We demonstrate that the expression of *Cdx* factors is necessary to establish spinal cord competency, by directly binding to neural regulatory regions that are accessible in hindbrain conditions (Figure 5D-F and TABLE S3) and by directly promoting posterior *Hox* genes (Figure S4E and TABLE S3). Hence, a major anterior-posterior division of the nervous system, separating the spinal cord from more rostral territories, is established prior to neural induction by the chromatin remodeling activities of CDX TFs.

Newly accessible genomic regions associated with the addition of WNT signals could be divided into those only transiently available and lost by the time cells had adopted a spinal cord identity (Figure 2A, NMP sites) and a set that continued to be accessible in spinal cord progenitors. This suggests in addition to ‘priming’ – identifying and making accessible – regulatory elements that are then sustained in spinal cord progenitors, CDX establishes a transition (“handover”) state between NMP and SC cells. This handover is driven by active remodeling of the chromatin both before (NMP sites; CDX-driven; Figure 5A) and after (SC sites; HOXC9/CDX-enriched; Figure 2J) neural induction. Moreover, CDX2 appears to repress directly genes involved in hindbrain neural identity (Figures 5E-F and S4E) indicating a pivotal function for CDX in securing spinal cord identity.

The importance of the CDX dependent genomic programme prior to neural induction is reinforced by a direct comparison of chromatin accessibility in hindbrain and spinal cord progenitors. In both conditions, cells have been exposed to RA and SHH signalling and express the neural progenitor transcription factors Sox1-3. Examining accessible genomic sites that distinguish hindbrain and spinal cord revealed an overrepresentation of the SOXB1 DNA binding motif in these sites. ChIP-seq confirmed that the binding site preference of SOX2 depends on AP position; hindbrain accessible regions, predicted to be bound by SOXB1 factors were enriched for SOX2 protein binding in hindbrain progenitors. Conversely, a different set of SOXB1 accessible sites, specific to the spinal cord, shows engagement of SOX2 at these sites in spinal cord but not hindbrain progenitors (Fig. 2). This suggests that deployment of SOX2 in neural progenitors is dependent on the genomic programme established in the epiblast from which the neural cells originate. The activity of CDX before neural induction directs neural TFs such as SOX2 to bind to spinal cord specific locations and abrogates binding to sites occupied only in the hindbrain (Figure 2I-J). CDX proteins regulate posterior *Hox* gene expression and establish the differences in *Hox* complement between spinal cord and hindbrain (Mazzoni et al., 2013; Nordström et al., 2006; van Rooijen et al., 2012b; Young et al., 2009). Thus, the differences in the *Hox* code between spinal cord and hindbrain appear responsible for the region-specific chromatin accessibility and distribution of SOX binding (Hagey et al., 2016). This highlights the importance of primary regionalisation in cells, preceding neural induction, to establish distinct hindbrain and spinal cord identities.

### Distinct lineages generate the anterior vs posterior nervous system

The divergence in the mechanisms for formation of the anterior and posterior nervous system is reminiscent of older ideas in which separate developmental organizers were proposed to induce different parts of the CNS (Mangold, 1933). This is consistent with experiments in chick embryos in which the identity of neural tissue induced by grafts of the organiser – tissue capable of inducing neural identity – depends on the embryonic age of the organiser: developmentally older organisers induce characteristics of the caudal neural tube but not forebrain (Storey et al., 1992). Thus, transplantation of different epiblast populations may have occurred at these different stages. In addition, it has been suggested that AP patterning events in the zebrafish epiblast is uncoupled from the neural inducing activities provided by the organizer (Koshida et al., 1998). Although in these studies definitive molecular markers were either lacking or specific to the hindbrain (but not spinal cord), these data hinted that axial patterning information might be established in the epiblast before the acquisition of neural identity. Indeed, NMPs, that generate both spinal cord and somites, are specified in the posterior epiblast in response to FGF and WNT signaling (Garriock et al., 2015; Tzouanacou et al., 2009; Wymeersch et al., 2016). Thus, spinal cord cells derive from a different lineage to the rest of the nervous system. These findings may reconcile old observations that grafts of tissue able to generate caudal structures frequently developed into both neural and mesodermal cell types, in contrast to the grafts producing forebrain structures that lacked mesodermal counterparts (Nieuwkoop, 1952).

The separate lineages generating hindbrain versus spinal cord progenitors prefigures differences in the complement of cell types generated in these regions. Serotonergic neurons and cranial visceral MNs are produced exclusively in the hindbrain (Carcagno et al., 2014; Cordes, 2001). This contrasts with preganglionic neurons of the sympathetic nervous system (Shirasaki and Pfaff, 2002) and the classes of somatic MNs and interneurons that are specific to the spinal cord (Jessell, 2000). Moreover, the genetic programme and progeny of neural crest derived from hindbrain and spinal cord levels differs (Simoes-Costa and Bronner, 2016); for example, the skeletogenic potential observed in cranial neural crest cells (NCCs), which form at hindbrain axial levels, is not observed in trunk NCCs generated at more posterior levels of the embryo (Martik and Bronner, 2017; Santagati and Rijli, 2003). Primary regionalisation events in the precursors that generate trunk, and not cranial, NCCs, may contribute to their inherent differences along the neuroaxis. Whether a similar strategy also accounts for the generation of posterior tissues contributing to mesodermal or endodermal lineages in the trunk of the embryo remains to be tested.

The transition from hindbrain to spinal cord specific cell types coincides with the approximate rostral limit of the neural progenitors lineage traced from *T/Bra* expressing NMPs (Garriock et al., 2015). This raises the possibility that the separate origins of the progenitors that populate the hindbrain and spinal cord underpins the regionally restricted generation of different neuronal subtypes. In this view, the differences in the ontogeny of the hindbrain and spinal cord establishes the distinct genomic regulatory environment responsible for generating region-specific cell types along the AP axis. This has implications for regenerative medicine and the design of conditions that produce neurons with authentic molecular identities. The finding that regionalisation is initiated and differences established prior to neural induction highlights the importance of determining the appropriate culture conditions at early stages of the directed differentiation of ESCs (Gouti et al., 2014; Lippmann et al., 2015; Maury et al., 2014). Focusing on these time windows and endeavouring to recapitulate normal developmental processes may contribute to deriving *in vitro* differentiated neuronal subtypes that accurately mimic their *in vivo* counterparts.

The role of the WNT-CDX genetic network in the specification of caudal tissue has been documented in a range of animals from across the bilaterian clade (Chawengsaksophak et al., 2004; Faas and Isaacs, 2009; Morales et al., 1996). The broad evolutionary conservation suggests that the network had this function in the last common ancestor of all Bilateria (Ryan et al., 2007). Whether the role of WNT-CDX activity in specifying distinct regions of the nervous system extends to non-vertebrates is yet to be firmly established. Nevertheless, the divergent lineage and distinct molecular events of the anterior and posterior nervous system is consistent with the proposed dual evolutionary origins of the central nervous system (Arendt et al., 2016). This hypothesis postulates that the bilaterian nervous system evolved from the merger of nerve centres residing at opposite poles of the ancestral pre-bilaterian animal (Arendt et al., 2016). In this view, the nervous system at the apical pole of the ancestral animal had a primary sensory function and modulated body physiology. Whereas the basally located blastoporal nervous system coordinated feeding movements and locomotion (Figure 6B). The expansion and fusion of these centres is proposed to have led to the bilaterian nerve cord and brain (Arendt et al., 2016; Tosches and Arendt, 2013). Hence, the distinct molecular mechanisms that specify anterior versus posterior vertebrate nervous systems may represent an evolutionary vestige of the processes that once generated neural tissue in pre-bilaterian animals.

## Methods

### Cell lines

All WT ESC culture was performed using the HM1 line (Doetschman et al., 1987). *Bra^-/-^* and *Cdx^1,2,4-/-^* knockout ESC lines were generated in the HM1 line using CRISPR as previously described (Gouti et al., 2017). Single guide RNAs were used to target the T-box domain *(T/Bra* mutant), and the caudal-like activation domain of *Cdx1, Cdx2* and *Cdx4* (*Cdx^1,2,4-/-^* triple mutant). Cell lines were validated by DNA sequencing and western blotting and routinely tested for mycoplasma.

### Cell culture and neural progenitor differentiation

Mouse ESCs were propagated on mitotically inactivated mouse embryonic fibroblasts (feeders) in DMEM knockout medium supplemented with 1000U/ml LIF (Chemicon), 10% cell-culture validated fetal bovine serum, penicillin/streptomycin, 2mM L-glutamine (Gibco). To obtain neural progenitors with anterior, hindbrain or posterior neural identity, ESCs were differentiated as previously described(Gouti et al., 2014). Briefly, ESCs were dissociated with 0.05% trypsin, and plated on gelatin-coated plates for two sequential 20 minute periods in ESC medium to separate them from their feeder layer cells which adhere to the plastic. To start the differentiation, cells remaining in the supernatant were pelleted by centrifugation, washed in PBS, and pelleted again. Cells were counted and resuspended in N2B27 medium containing 10ng/ml bFGF to a concentration of 10^6^ cells per ml, and 50,000 cells per 35mm CELLBIND dish (Corning) were plated. N2B27 medium contained a 1:1 ratio of Advanced Dulbecco’s Modified Eagle Medium F12:Neurobasal medium (Gibco) supplemented with 1xN2 (Gibco), 1xB27 (Gibco), 2mM L-glutamine (Gibco), 40 μg/ml BSA (Sigma), penicillin/streptomycin and 0.1mM 2-mercaptoethanol. To generate anterior neural progenitors, the cells were grown up to day 3 in N2B27 + 10ng/ml bFGF, followed by N2B27 + 500nM smoothened agonist (SAG; Calbiochem) from day 3-5. To generate hindbrain neural progenitors, cells were cultured under the same conditions as the anterior, but were additionally exposed to 100nM retinoic acid (RA; Sigma) from day 3-5. To generate spinal cord neural progenitors, cells were cultured with N2B27 + 10ng/ml bFGF until day 2, N2B27 + 10ng/ml bFGF + 5μM CHIR99021 (Axon) until day 3, and N2B27 + 100nM RA + 500nM SAG until day 5. For Hindbrain+ treated cells (Fig. 3), cells were differentiated under hindbrain conditions with one modification between day 4-5, where they were additionally exposed to 10ng/ml bFGF and 5μM CHIR99021 in addition to continued treatment with RA and SAG as above. For all differentiations, media changes were made every 24 hours from day 2.

### Immunofluorescence and microscopy

Cells were washed in PBS and fixed in 4% paraformaldehyde in PBS for 15min at 4 degrees, followed by two washes in PBS and one wash in PBST (0.1% Triton X-100 diluted in PBS). Primary antibodies were applied overnight at 4 degrees diluted in filter-sterilized blocking solution (3% FBS diluted in PBST). Cells were washed 3x in PBST and secondary antibodies (AlexaFluor conjugated; Invitrogen) were applied at room temperature, diluted 1:1000 in PBS for 1 hr. Cells were washed 3x in PBS, incubated with DAPI for 5 min in PBS and washed twice before mounting with Prolong Gold (Invitrogen). Primary antibodies were diluted as follows: Phox2b rabbit, kindly provided by Jean-Francois Brunet (Pattyn et al., 1997), 1/1000; Olig2 rabbit (Millipore, AB9610; 1/1000); Olig2 guinea-pig, kindly provided by Ben Novitch (Novitch et al., 2001), 1/1000, and Sox2 goat (R&D Systems, AF2018; 1/500). Cells were imaged on a Zeiss Imager.Z2 microscope using the ApoTome.2 structured illumination platform. Z-stacks were acquired and represented as maximum intensity projections using ImageJ software. Immunofluorescence was performed on a minimum of 3 biological replicates, from independent experiments.

### RNA extraction, cDNA synthesis and qPCR analysis

RNA was extracted from cells using a Qiagen RNeasy kit, following the manufacturer's instructions. Extracts were digested with DNase I to eliminate genomic DNA. First strand cDNA synthesis was performed using Superscript III (Invitrogen) using random hexamers and was amplified using Platinum SYBR-Green (Invitrogen). qPCR was performed using the Applied Biosystems 7900HT Fast Real Time PCR. PCR primers were designed using NCBI primer blast or primer3 software, using exon-spanning junctions where possible. Expression values for each gene were normalised against ß-actin, using the delta-delta CT method. Error bars represent standard deviation across three biological replicate samples. qPCR was performed on 3 biological replicates for every primer pair analysed. Primer sequences are available in Table 4.

### ATAC-seq

ATAC-seq was performed on ESCs and at each day of the differentiation following methods previously described (Buenrostro et al., 2013, 2015). Briefly, adherent cells were treated with StemPro Accutase (A1110501) to obtain a single cell suspension. Cells were counted and resuspended to obtain 50,000 cells per sample in ice-cold PBS. Cells were pelleted and resuspended in lysis buffer (10mM Tris-HCl pH 7.4, 10mM NaCl, 3mM MgCl2, 0.1% IGEPAL). Following a 10 min centrifugation at 4°C, nucleic extracts were resupsended in transposition buffer for 30 minutes at 37°C and purified using a Qiagen MinElute PCR Purification kit following manufacturer’s instructions. Transposed DNA was eluted in a 10μl volume and amplified by PCR with Nextera primers (Buenrostro et al., 2013) to generate single-indexed libraries. A maximum of 12 cycles of PCR (determined using optimisation experiments) was used to prevent saturation biases based on optimisation of qPCR cycles as previously described (Buenrostro et al., 2015). Library quality control was carried out using the Bioanalyzer High-Sensitivity DNA analysis kit. Libraries were sequenced as paired-end 50 or 100 bp reads, multiplexing 4 samples per lane on the Illumina High-Seq 2500 platform at the Francis Crick Institute Advanced Sequencing Facility. For all conditions, two biological replicate samples were collected from independent experiments.

### *In vivo* ATAC-seq and mouse lines

*Sox2eGFP* heterozygous mice (Ellis et al., 2004) were maintained on a C57BL6 background. To obtain embryos, *Sox2eGFP* heterozygous mice were mated to wild type mice. Embryos for ATAC-seq were harvested at e9.5 in HBSS buffer (GIBCO) containing 5% FBS. As the ratio of cells to transposase is a critical parameter in generating ATAC-seq results (Buenrostro et al., 2015), we aimed to use the same ratio of cells *in vivo* as in vitro for maximum comparability. To obtain sufficient quantities of cells from *in vivo*, embryos from several litters were pooled together and screened for GFP using a Leica MZFL widefield microscope with a GFP filter set. Embryos were separated into GFP positive and negative pools. To enrich for anterior and spinal cord neural progenitors, GFP positive embryos were dissected as follows: heads were decapitated at the second pharyngeal arch and otocysts removed to avoid contamination with other GFP-expressing cells. To obtain spinal cord NPs, the neural tube and surrounding somitic tissue was dissected, from the level of caudal hindbrain to the tailbud posterior neuropore. Both cranial and trunk regions were minced with forceps, incubated for 5 minutes on ice in enzyme-free dissociation buffer (Gibco) and then gently passed through a 40μm filter using the plunger from a sterile syringe. Dissociated cells were collected, centrifuged at 4°C for 5 minutes at 1500rpm and resuspended in 500ul HBSS buffer containing 5% FBS. Cells were passed through a 40μm filter and sorted using flow cytometry. Flow analysis and sorting was performed by the Francis Crick Flow Cytometry facility, using an Aria Fusion cell sorter with a 488nm laser. GFP negative cells (obtained from negative littermates collected in parallel) were used as a negative control to set voltage gating. 50,000 GFP positive cells from anterior and spinal cord levels obtained from FACS were subject to ATAC-seq as described for in vitro-derived cells. Duplicate samples were collected on independent days to represent biological repeats. All animal procedures were performed in accordance with the Animal (Scientific Procedures) Act 1986 under the UK Home Office project licences PPL80/2528 and PD415DD17.

### ChIP-seq

Sox2 ChIP-seq (Santa cruz antibody SC-17320X) was performed in duplicate as previously described (Kutejova et al., 2016). Briefly, 10-30 million cells from day 5 hindbrain or day 5 spinal cord neural progenitors were crosslinked in 1% formaldehyde for 20 min at 4 degrees. Chromatin was sonicated using a Diagenode Bioruptor (using a cycle of 30sec on, 30 sec off) until fragments were between 200-400bp. 3ug antibody was incubated together with cell lysate overnight at 4°C on a rotating wheel. Immunoprecipitation of chromatin fragments were captured using Protein G-coupled Dynabeads (Life Technologies). Samples were decrosslinked and purified using the Qiagen MinElute kit. Approximately 10ng ChIP DNA and 10ng input DNA for each condition was used to prepare ChIP-seq libraries using the KAPA Hyper Prep Kit (Illumina). Biological duplicates were obtained for both conditions from separate experiments. Libraries were sequenced as single-end, 50bp reads on the Illumina High-Seq 2500 platform (Francis Crick Institute).

### Data analysis

**ATAC-seq pre-processing**

Sequencing adapters and poor quality bases were trimmed from reads using *trim_galore* with default settings (https://www.bioinformatics.babraham.ac.uk/projects/trimgalore/). Reads were aligned to the *mm10* reference genome using *bowtie2* with parameters: *-X 2000* --- *sensitive-local* (Langmead and Salzberg, 2012). Alignments were filtered for unmapped, low quality (MAPQ < 30), mapping to chrM or not properly paired fragments. PCR duplicates were marked using *Piccard* and not used in subsequent analysis steps.

Signal tracks were computed as *fragments per million per base pair* (FPM) using *deepTools bamCoverage* with following settings: *--scaleFactor 10^6^/Library size -bs 1 -extendReads -samFlagInclude 66 -ignoreDuplicates* (Ramírez et al., 2016). Read enriched regions were called with MACS2 using the options *-g mm -p 0.01 - nomodel-f BAMPE* (Zhang et al., 2008).

For analysis on insertion level, such as TF footprinting, we adjusted plus-strand insertions by +4bp and minus-strand insertions by -5bp (Buenrostro et al., 2013).

#### ATAC-seq - differential and clustering analysis

Initially, a robust peak set per condition was obtained by computing the irreproducible discovery rate (IDR) between duplicates and thresholding for peaks with an IDR <= 0.1 (Li et al., 2011). These robust peak sets were merged to one consensus region set using bedtools merge (Quinlan and Hall, 2010). We computed the fragment coverage for all samples of these regions using featureCounts with additional options –F SAF –p –ignoreDup (Liao et al., 2013).

The resulting count table was used as input for a differential chromatin accessibility analysis with DESeq2 (default settings) to compare pairwise all WT in vitro conditions with D0 (ESC) (Love et al., 2014). We obtained a set of variable regions by filtering for differential peaks with log2(FC) > 1 and adjusted p-value < 0.01.

These variable regions were clustered in order to resolve the complex chromatin dynamics from our multiple condition time series data. For this, DESeq2 normalized count data was transformed by computing region-wise z-scores (z=(x – mean(x))/sd(x)). Variable regions were clustered using the z-score matrix as input for a self-organizing map (SOM) with a grid of 5x5 cells, a hexagonal topology and a Gaussian neighborhood function. We found that 5x5 best reflects the complex chromatin dynamics, though different grid sizes largely reproduced the same dynamics. SOM clusters were merged by similar trends and manually annotated using known regulatory regions.

#### ATAC-seq - peak annotation

Consensus regions were annotated with nearby genes using *ChIPseeker annotatePeak* with GENCODE release M14 (Mudge and Harrow, 2015). Regions were further annotated to nearby genes using default settings (basal rule) in GREAT (McLean et al., 2010). Each gene is assigned to a regulatory region spanning 5kb upstream and 1 kb downstream of the TSS (irrespective of other genes). This regulatory domain is extended in both directions to all nearest genes, up to a maximum of 1000kb (McLean et al., 2010).

#### ATAC-seq – *In vitro* vs *in vivo* comparative analysis

Fragment coverage of *in vitro* consensus regions for *in vivo* ATAC-seq experiments was computed using *featureCounts* with the same options as above. We normalized counts from *in vivo* and corresponding *in vitro* samples using DESeq2 with default settings. The normalized counts were used to compute log2 fold-changes between conditions for *in vivo* and *in vitro* respectively. These were filtered for regions falling into the SOM cluster: Anterior, Neural and Spinal cord. Distribution of log2 fold-changes were compared with two-sided Wilcoxon rank sum tests. ATAC-seq meta profiles were plotted for regions which were enriched in both *in vitro* and the corresponding *in vivo* sample (log2(FC) > 0.5).

#### ATAC-seq - ENCODE DHS overlap

DNAse hypersensitive sites (DHS) for various tissues, on the mm10 reference genome, were obtained from the ENCODE data portal (Consortium, 2012; Sloan et al., 2016). DHS regions were overlapped with our SOM clustered regions using *GenomicRanges findOverlaps* (Lawrence et al., 2013).

#### ATAC-seq - Vista enhancer enrichment

Vista enhancer regions were downloaded from the ENCODE data portal (Consortium, 2012; Sloan et al., 2016) (https://www.encodeproject.org/). Enhancers were overlapped with our SOM clustered regions and enrichment was examined with a one-sided binomial test, similar to methods used for GREAT analysis (McLean et al., 2010). Resulting p-values were corrected for multiple testing using the Benjamini-Hochberg procedure.

#### ATAC-seq - motif enrichment

Regions were centered on the summit, rescaled to 300bp using *bedtools slop* and sequences were queried by *bedtools* getfasta from the mm10 reference genome. Resulting fasta files were used as input for *HOMER findMotifs.pl* with parameters – *bits-mset vertebrates* (Heinz et al., 2010). TF motifs were visualized using ggSeqLogo (Wagih, 2017).

#### ATAC-seq - TF footprinting

TF binding motifs for factors of interest were queried from the JASPAR database (Khan et al., 2017).We matched these motifs against genomic sequences using motifmatchr to obtain their genomic positions (https://github.com/GreenleafLab/motifmatchr). Two strategies were used depending on the question: either the motif was genome-wide matched (for CTCF; Figure S1F) or +/-5kbp around peak summits was used (for CDX; Figure S4D).

Resulting motif positions were extended +/-150bp. Adjusted Tn5 insertions from fragments <= 100bp were counted per base-pair and strand. For the footprinting, we used PWM scores and corresponding insertion count matrix as input for CENTIPEDE to compute posterior probabilities that a motif is bound (Pique-Regi et al., 2011). A different threshold to classify bound/unbound was used depending on the motif matching strategy (genome-wide: >= 0.99; peak summit: >= 0.9).

#### ChIP-seq pre-processing

Sequencing adapters and poor quality base calls were trimmed from reads using *trim_galore* with default settings. Trimmed reads were aligned against the mm10 reference genome using *bowtie2* with --*sensitive* as additional option. Alignments were filtered for unmapped, multi-mapping and duplicated reads.

Signal tracks as log2 fold-change between ChIP and input were generated using *deepTools bamCompare* with following parameters --*scaleFactorsMethod SES-ratio log2-bs 25-ignoreDuplicates* (Ramirez et al., 2016).

Peak calling was performed using MACS2 with --g *mm --p 0.001* (Zhang et al., 2008). Re-analysis of publicly available datasets was performed in the same way as for new samples. All ChIP-seq datasets used in this study are listed in Table S2.

#### ChIP-seq enrichment analysis

Enrichment analysis of TF peaks was performed with LOLA (default settings) using the set of variable ATAC-seq regions as universe (Sheffield and Bock, 2015). We considered all TF peak sets with an adjusted p-value < 0.01 as enriched.

To complement the mm10 core database with TFs relevant for neural development, we added 4 new samples and 39 publicly available TF ChIP-seq datasets (Table S2). Replicates of ChIP-seq experiments were considered separately where available.

#### RNA-seq pre-processing

RNA-seq experiments in this study were quantified using *Salmon* (quasi-mapping mode) with the GENCODE release M14 (Mudge and Harrow, 2015). Single-end as well as paired-end reads were processed using following options: -*I A --seqBias - numBootstraps 50*. Resulting counts and transcripts per million (TPM) were used for downstream analysis.

Differential analysis of D5H vs D5SC (Gouti et al., 2014) was performed using DESeq2 with default settings. Resulting adjusted p-values were used for Figure S4E.

#### Interaction database

A dataset of putative gene-region interactions was downloaded from the 4DGenome database (https://4dgenome.research.chop.edu/). The obtained interactions were mapped to *mm9* and for further downstream analysis re-mapped to *mm10* using *UCSC-liftOver* with default setting. Interactions for which only one anchor could be mapped to *mm10* were removed.

Putative chromatin-chromatin interactions were mapped by filtering for anchors which overlap open chromatin sites from this study.

#### Gene ontology enrichment of CDX2 bound open chromatin sites

MNP CDX2 ChIP-seq peaks from Mazzoni et al., 2013, were overlapped with NMP, NMP-SC, spinal cord, H/SC and hindbrain regions. Resulting peak sets were used as input for gene ontology enrichment analysis using GREAT (default settings) (McLean et al., 2010).

## Data availability

All data generated in this study is available from the Array Express website.

## Code availability

Analysis scripts are available at https://github.com/luslab/NeuralATACseq

## Author contributions

VM and JB conceived the project, interpreted the data and wrote the manuscript. VM designed and performed the experiments and performed data analysis. SS and EP developed custom code and performed data analysis. DS assisted in the collection of *in vivo* samples and constructed the *T/Bra* and *Cdx* mutant lines. MG performed the initial differentiations with the *T/Bra* and *Cdx* mutant lines and critically revised the manuscript. RLB provided access to animal colonies and performed critical revision of the manuscript. NML interpreted the data and critically revised the manuscript.

## Acknowledgements

The authors would like to thank the Science Technology Platforms at the Francis Crick Institute (and formerly at the National Institute for Medical Research). In particular, we thank the Advanced Sequencing Facility, Bioinformatics And BioStatistics Facility, the Flow Cytometry Facility, High Performance Computing and the Biological Research Facility for their ongoing support and access to equipment. We are very grateful to Karine Rizzoti for providing *Sox2eGFP* mice, Jean-Francois Brunet for the Phox2b antibody and Bennett Novitch for the Olig2 antibody. The authors also acknowledge and thank the ENCODE Consortium and the production labs of John Stamatoyannopoulos (UW) and Ross Hardison (PennState). We thank Alfonso Martinez-Arias, Francois Guillemot, Jens Kleinjung and Karine Rizzoti for comments on the manuscript and members of the Briscoe lab for useful discussion. We thank Boris Lenhard, Julien Delile and Jens Kleinjung for critical feedback on data analysis. This work was supported by the Francis Crick Institute, which receives its core funding from Cancer Research UK (FC001051, FC0010110), the UK Medical Research Council (FC001051, FC0010110), and the Wellcome Trust (FC001051, FC0010110); JB and NML are funded by the Wellcome Trust (WT098326MA); JB by the BBSRC (BB/J015539/1).

## SUPPLEMENTARY FIGURE LEGENDS

**Figure S1.**
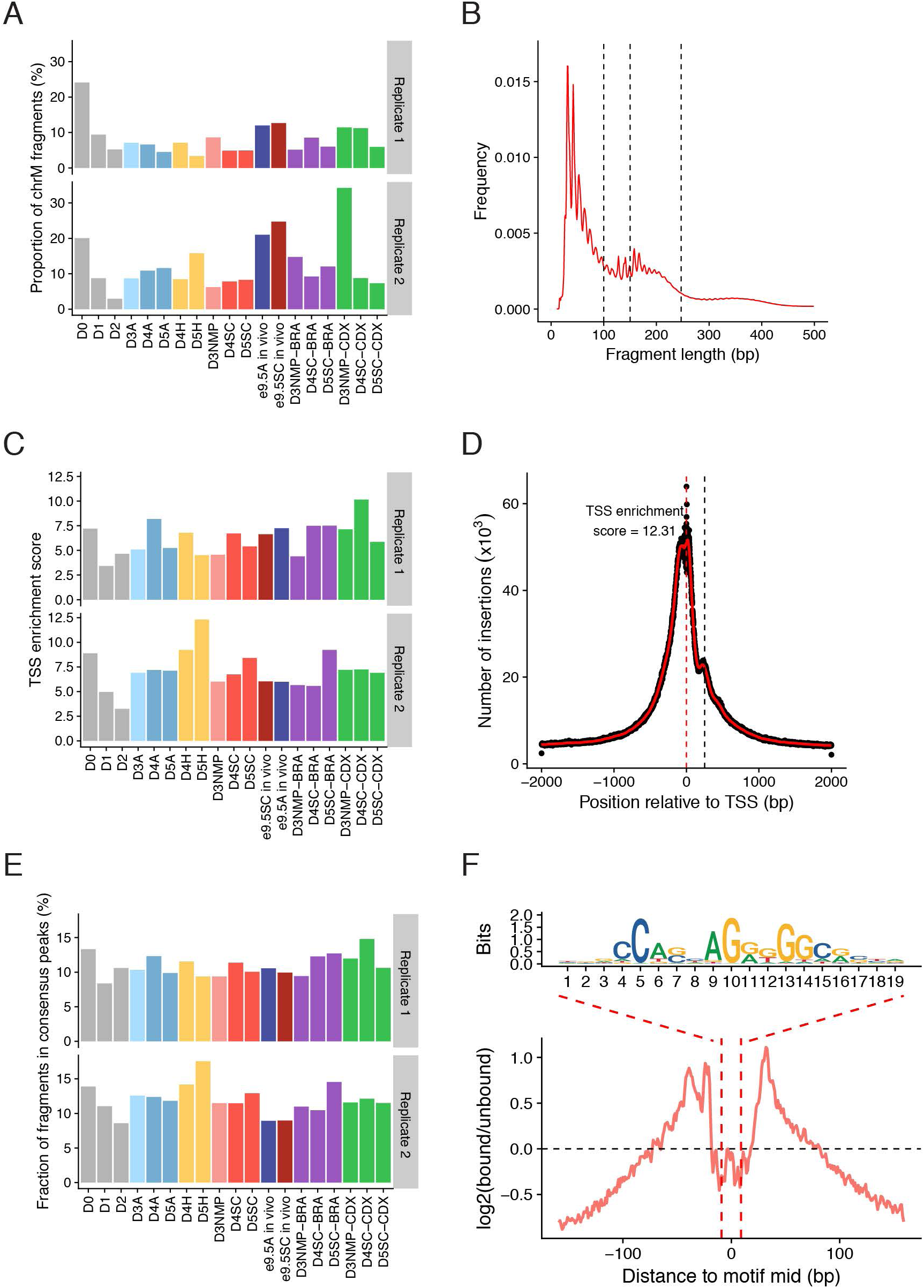
Quality control for all ATAC-seq samples generated in this study. **(A)**The proportion of mitochondrial fragments recovered across each sample. **(B)** Representative example showing the distribution of fragment lengths recovered from ATAC-seq, using paired-end sequencing. **(C)** Average level of Tn5 enrichment (score = maximum(number of insertions)/minimum(number of insertions)) observed across transcription start sites (TSS) for each sample. **(D)** Summarised Tn5 insertion profile covering +/-2kb around annotated TSS for sample D5H (replicate 2). Red line corresponds to a 50bp running average. **(E)** The fractions of fragments that map to *in vitro* consensus peak regions. **(F)** CTCF footprint present in ESC accessible regions as determined by ATAC-seq.

**Figure S2.**
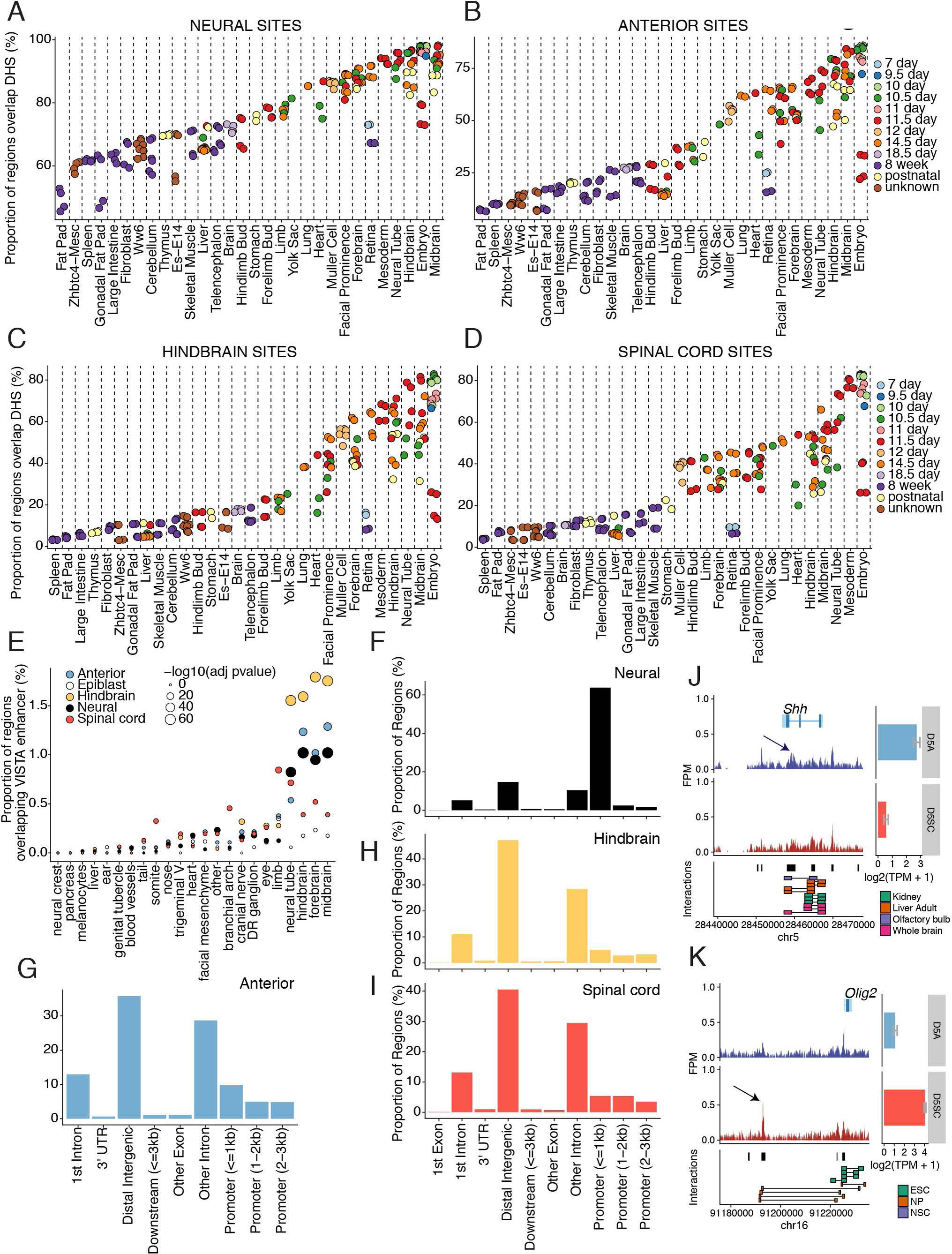
Tissue specificity and genomic location of regulatory regions that define neural and region-specific identity. **(A-D)** Comparison of ATAC-seq identified regions with previously published DNase hypersensitivity sites present across a range of *in vivo* tissues and time points from the ENCODE regulatory element database (Consortium, 2012; Sloan et al., 2016). Genomic regions correspond to neural (A), anterior (B), hindbrain (C) and spinal cord (D) specific sites from Figure 2A. Each set of genomic regions demonstrates an enrichment in embryonic and neural samples *in vivo*. **(E)** Comparison of ATAC-seq identified regions with the Vista enhancer database (Visel et al., 2007) shows that ATAC-seq recovers enhancers that show neural tissue specificity *in vivo*. **(F-I)** Classification of neural (F), anterior (G), hindbrain (H) and spinal cord (I) sites according to genomic position. Neural sites are enriched at promoter regions (F), in contrast to the region-specific sites, which predominantly occupy distal intergenic and intronic regions (G-I). **(J-K)** Genome browser view (mm10 assembly) showing ATAC-seq from anterior (blue track) and spinal cord (red track) neural progenitors obtained from e9.5 mouse embryos at the *Shh* (J) and *Olig2* (K) locus. Arrows indicate known enhancers that direct *Shh* expression in the midbrain (Epstein et al, 1999; J) and *Olig2* in the spinal cord (Oosterveen et al, 2012 and Peterson et al, 2012; K). Gene expression levels determined by mRNA-seq (Gouti et al., 2014) are shown as bar plots from *in vitro* Day 5 anterior (blue) and spinal cord (red) conditions (error bars = SEM). Chromatin interactions recovered from indicated tissues are presented below for comparison. Peak regions are represented with black bars. A=anterior neural progenitor; DR=dorsal root; NP=neural progenitor; NSC=neural stem cell; SC=spinal cord progenitor; TPM=transcripts per million.

**Figure S3.**
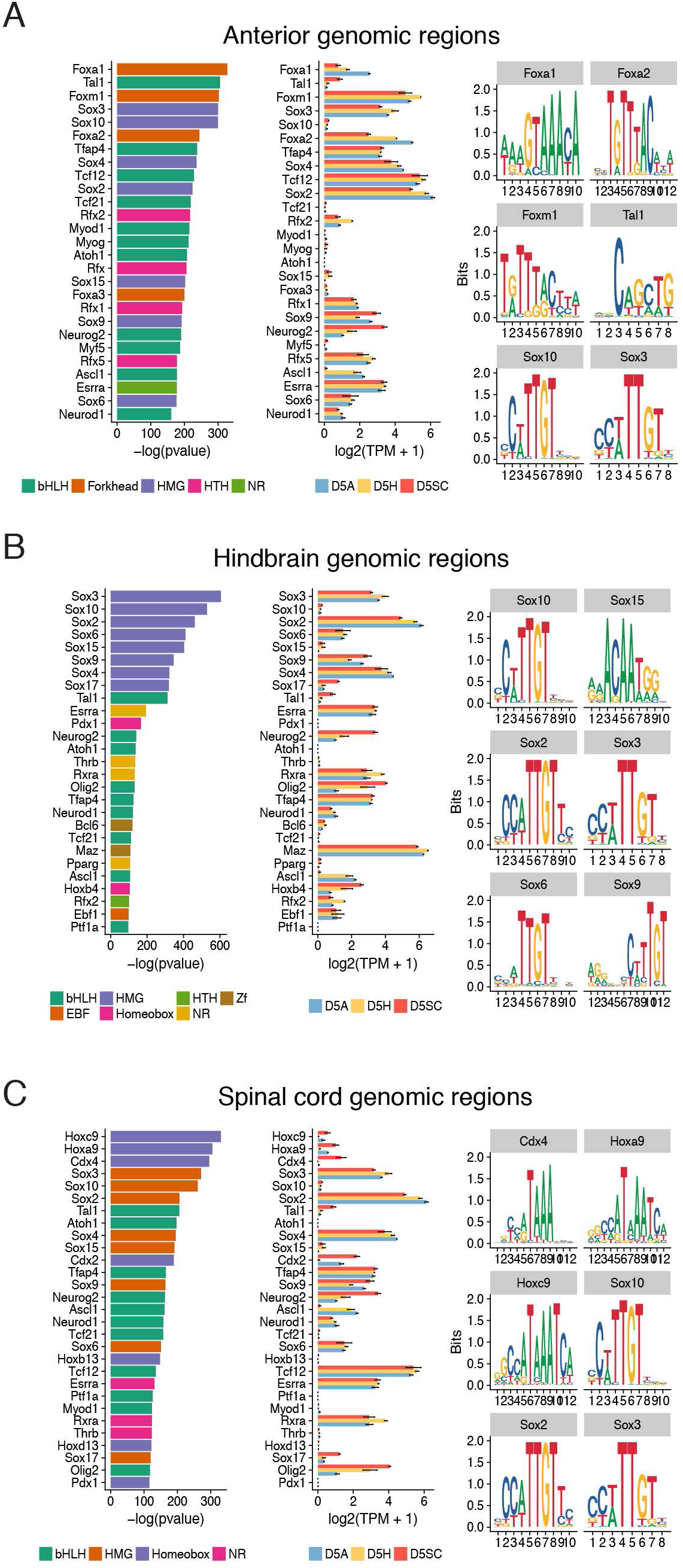
Motif analysis of region specific sites that define anterior, hindbrain and spinal cord. **(A-C)** Motif analysis performed using Homer (Heinz et al., 2010) on anterior (A), hindbrain **(B)** and spinal cord (**C)** specific sites shows distinct and common neural factors are enriched at each AP level. For each predicted factor (shown on the left), their corresponding expression level determined by mRNA-seq (Gouti et al., 2014) in the same condition at D5 of the *in vitro* differentiation is shown (central column; error bars = SEM). The top 6 predicted motif logos are presented on the right. TPM=transcripts per million.

**Figure S4.**
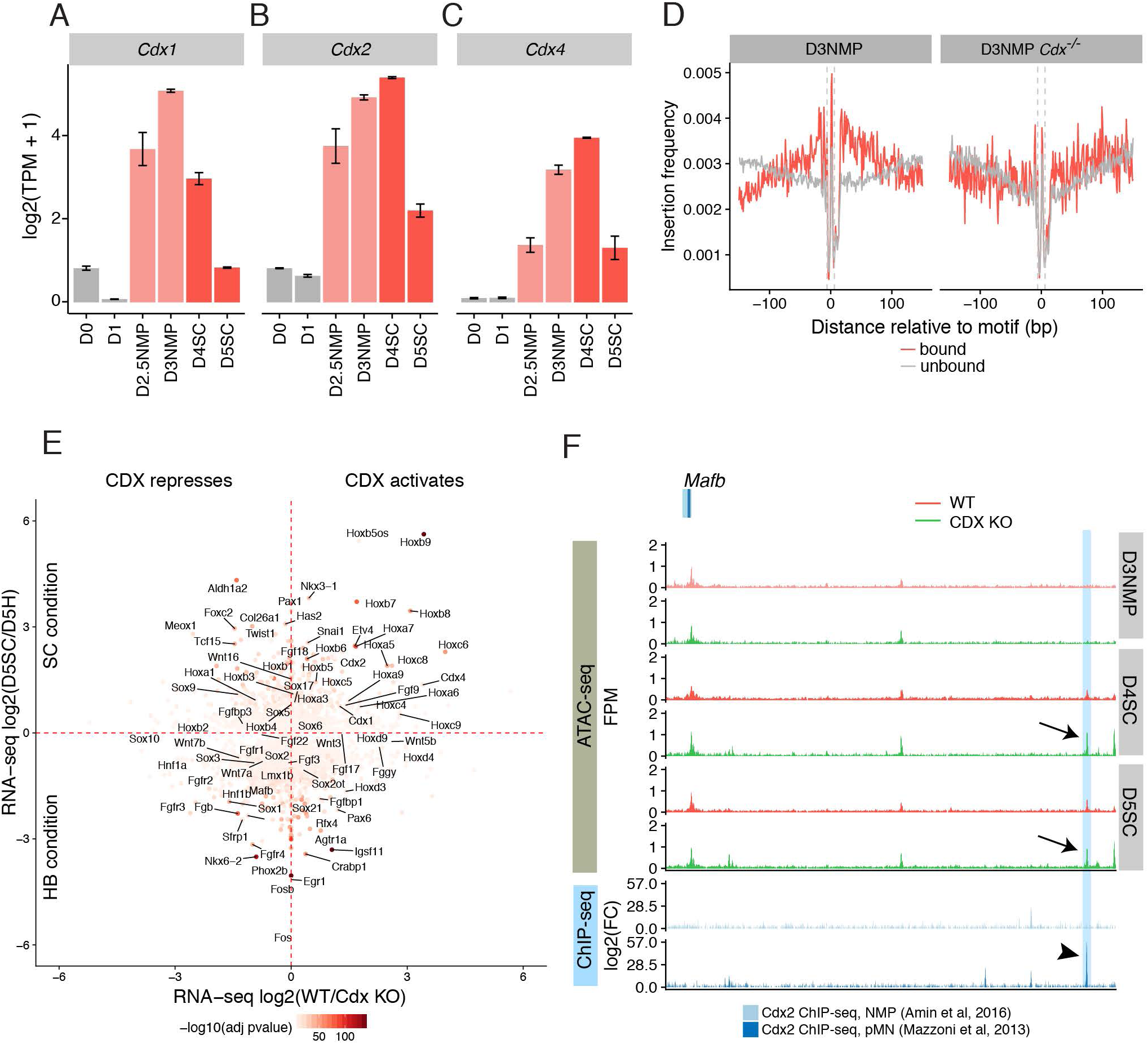
Expression dynamics of *Cdx* TFs during the spinal cord differentiation. **(A-C)** Expression profile determined by mRNA-seq for *Cdx1* (A), *Cdx2* (B) and *Cdx4* (C) from day 0 to day 5 of the spinal cord differentiation. **(D)** Nucleotide resolution of the frequency of integration of the Tn5 transposon reveals increased engagement of CDX2 at NMP accessible sites in WT compared to *Cdx* mutant D3NMP sites. **(E)** Differential gene expression determined by mRNA-seq in Day 5 hindbrain (D5HB) versus Day 5 spinal cord (D5SC) *in vitro* conditions (Gouti et al., 2014) compared with wildtype (WT) and *Cdx* mutant *(Cdx* KO) *in vivo* samples from microdissected e8.0 posterior tailbud tissue (Amin et al, 2016). CDX positively regulates *Hoxb9* and other 5’ *Hox* genes while it represses *Aldh1a2* in the spinal cord, in agreement with previous studies (Gouti et al., 2017). CDX negatively regulates many hindbrain genes including *Mafb*. **(F)** ATAC-seq from wildtype (red) and *Cdx* mutant (green) cells at indicated stages (grey bars) at the *Mafb* genomic region. Arrows indicate ectopic accessibility observed in *CDX* mutant cells between Day 4-5 of the spinal cord differentiation. This region (blue shading) overlaps with a binding site occupied by CDX in motor neuron progenitor (pMN) conditions (arrowhead) from previously published studies (Mazzoni et al., 2013). NMP=neuromesodermal progenitor; SC=spinal cord; TPM=transcripts per million. Error bars = SEM.

**Figure S5.**
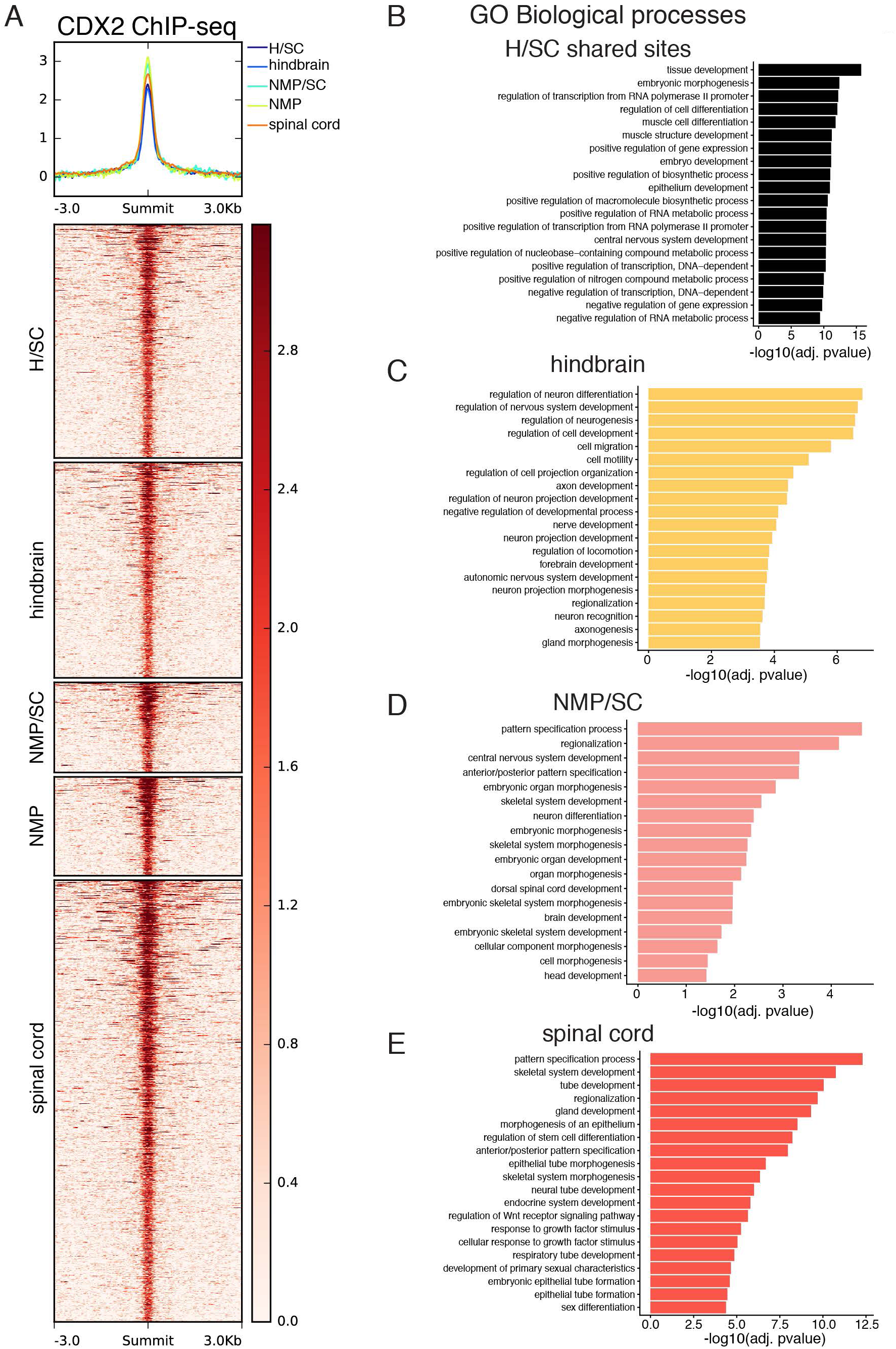
CDX2 occupancy in open chromatin sites and associated gene ontology enrichment. **(A)** Region heatmap displays CDX2 ChIP-seq binding at open chromatin sites recovered from the self-organising map (pMN; Mazzoni et al., 2013). **(B-E)** Gene ontology enrichment analysis for CDX2-bound regions shown in (A). In hindbrain accessible regions (C), CDX2 binding is associated with neural genes in contrast to either the NMP and spinal cord (NMP/SC) shared or SC-specific sites (E), which target genes involved in anterior-posterior patterning.

## Supplementary Tables

**Table S1**. Peak to gene annotation of region-specific sites identified in this study.

**Table S2.** List of all datasets used for ChIP-seq, ATAC-seq, and mRNA-seq analysis.

**Table S3.** List of NMP sites, NMP/SC shared and SC sites.

**Table S4**. Primers used for qPCR.

